# Virulent disease epidemics can increase host density by depressing foraging of hosts

**DOI:** 10.1101/2020.07.06.189878

**Authors:** Rachel M. Penczykowski, Spencer R. Hall, Marta S. Shocket, Jessica Housley Ochs, Brian C. P. Lemanski, Hema Sundar, Meghan A. Duffy

**Affiliations:** School of Biology, Georgia Institute of Technology, Atlanta, GA 30332 USA; Department of Biology, Washington University in St. Louis, St. Louis, Missouri 63130 USA; Department of Biology, Indiana University, Bloomington, IN 47405 USA; Department of Biology, Colgate University, Hamilton, NY 13346 USA; Department of Ecology and Evolutionary Biology, University of Michigan, Ann Arbor, MI 48109 USA

**Keywords:** foraging depression, host-parasite, host-resources, hydra effect, trait-mediated indirect effect, density-mediated indirect effect, compensatory population growth, illness-mediated anorexia

## Abstract

All else equal, parasites that harm host fitness should depress densities of their hosts. However, parasites that alter host traits may increase host density via indirect ecological interactions. Here, we show how depression of infected host foraging rate can produce such a hydra effect. Using a foraging assay, we quantified reduced foraging rates of a zooplankton host infected with a virulent fungal parasite. We then parameterized a dynamical model of hosts, parasites, and resources with this foraging function, showing how foraging depression can create a hydra effect. Mathematically, the hydra arose when increased resource productivity exceeded any increase in resource consumption per host. Therefore, the foraging-mediated hydra effect more likely emerged (1) for hosts which strongly control logistic-like resources and (2) during larger epidemics of moderately virulent parasites. We then analyzed epidemics from 13 fungal epidemics in nature. We found evidence for a foraging-mediated hydra effect: large outbreaks depressed foraging rate and correlated with increased densities of both algae and hosts. Therefore, depression of foraging rate of infected hosts can produce higher host densities even during epidemics of parasites that increase host mortality. Such hydras might prevent collapse of host populations but also could produce higher densities of infected hosts.

## Introduction

Disease epidemics can drive declines in host populations (Anagnostakis 1982; Daszak et al. 1999; Frick et al. 2010; Lessios et al. 1984), trigger conservation crises for wildlife such as mammals (Roelke-Parker et al. 1996) and birds (Cooper et al. 2009; Hochachka and Dhondt 2000), and even sometimes drive hosts extinct (amphibians: (Vredenburg et al. 2010)). Disease outbreaks can also damage economically valuable crops (Fry and Goodwin 1997) and livestock (Cleaveland et al. 2001). Even worse, climate change can further exacerbate disease epidemics (Altizer et al. 2013; Sanderson and Alexander 2020; Shocket et al. 2018). Therefore, it is imperative to identify when, where, and why parasites depress density of their hosts during epidemics.

Typically, we predict that parasites depress host density because infection exacts virulent costs to host fitness. Indeed, infection often can increase mortality rate and/or decrease fecundity of infected hosts. Simple disease models illustrate how those two factors can lower host density relative to disease-free conditions (Anderson and May 1979; Anderson and May 1981). Furthermore, that harm can become amplified by higher transmission of disease (which can lead to higher prevalence of infection). Higher transmission results from higher per capita exposure and/or susceptibility (the product of which is called ‘transmission rate’ (Dwyer and Elkinton 1993; Strauss et al. 2018)). Additionally, higher transmission can occur in more enriched systems that support higher density of hosts (assuming density-dependent spread of disease: (Johnson et al. 2010)). Therefore, we might expect larger absolute and/or relative depression of host density when virulent parasites reach higher prevalence.

On the other hand, this above outcome might reverse when infection depresses foraging rate of hosts. Many parasites lower foraging rate of hosts (Hite and Cressler 2019; Hite et al. 2020; Strauss et al. 2019). At first glance, such foraging depression — whether a defense strategy or fitness cost of infection (Hite et al. 2020) — might seem to exacerbate declines of host density during epidemics. After all, lower intake of energy, when coupled with reduced survivorship and/or fecundity from infection, might harm fitness of hosts even more (all else equal). However, we show that foraging depression can sometimes *increase* host density through a hydra effect (Abrams 2009). This outcome requires that hosts must strongly control a dynamic resource, and that the resource must reach highest productivity at intermediate density (e.g., growing logistically or logistic-like). When those conditions are met, foraging reduction can increase density, and hence production, of resources through indirect feedbacks. Those increases in resource production can then compensate for increased energetic demands of hosts (a consequence of virulence). When the shift in resource production exceeds that in resource consumption, a foraging-mediated hydra effect emerges, leading to higher host density in the presence of parasites — even during very large outbreaks.

Here, we illustrate this foraging-mediated hydra mechanism using a freshwater system with a zooplankton host (*Daphnia dentifera*). This host can strongly depress an algal resource that reaches highest production at intermediate density. This host becomes infected by a fungus that predominantly lowers survival (Hall et al. 2010) rather than fecundity (unlike, e.g., the bacterium *Pasteuria*: (Auld et al. 2012)). In this study, we demonstrate that infection also lowers foraging rate of hosts in an experiment, particularly as the final transmission stage of the fungus (spores) accumulate within the body cavity. We then parameterized a foraging depression function and incorporated it into a dynamical model. The model revealed how epidemics can drive higher host density (similar to how predators can increase prey density through changes in foraging behavior: (Peacor and Werner 2001)). This foraging-mediated hydra effect becomes more likely as epidemics become larger (e.g., with higher accrual of spores within hosts and higher carrying capacity of the resource) and with stronger foraging depression. Conversely, it becomes less likely with higher virulence on survival. Finally, a survey of fungal epidemics in lakes showed that larger epidemics (with greater infection prevalence) yielded higher parasite production per host. We estimated the depression in foraging due to disease in those lakes, and found that lower foraging correlated with joint increases of algal and zooplankton populations during epidemics. Taken together, this combination of experiments, dynamical modeling, and field surveys demonstrates how foraging depression can increase host density during epidemics of parasites that kill their hosts.

### Study System and a Function for Foraging Depression (a model competition)

#### Disease system

The focal host, the zooplankton *Daphnia dentifera*, strongly grazes on phytoplankton in many lakes throughout the Midwestern USA (Tessier and Woodruff 2002). Hosts ingest infectious propagules (spores) of the parasitic fungus *Metschnikowia bicuspidata* while foraging on small (< 80 µm) phytoplankton (Hall et al. 2007b). As the parasite fills its host’s hemolymph with spores (Ebert 2005; Green 1974), it reduces host growth, fecundity, and survivorship (Hall et al. 2009b). Death of the infected host releases spores into the water to then infect new hosts. Sometimes, epidemics of this fungus reduce host density and indirectly increase density of the algal resource via a trophic cascade (Duffy 2007; Hall et al. 2011). At other times, host density remains high during epidemics (Duffy and Hall 2008; Hall et al. 2011).

#### Foraging rate experiment: methods

We estimated foraging rate using an experiment, summarized only briefly here (see Appendix Section 1 for details). We measured feeding rate on individuals of cohorts of uninfected and infected hosts. To create a gradient in body size (host length, *L*_*H*_) and spore accumulation (*σ*), we measured food consumption by individuals of progressively older (and for the infected class, more infected) age cohorts. Thus, we placed individual hosts into small tubes containing their algal food, allowed them to graze for a short period of time (four hours), measured remaining food using a fluorimeter, and estimated length using a dissecting microscope. We ensured infection status by smashing hosts to release spores contained in their body. These spores represent the final life stage of the parasite; their presence indicates terminal infection (Stewart Merrill et al. 2019).

#### Functions for foraging depression: candidate models

We statistically competed models linking spore accumulation (*σ*) and body size (host length, *L*_*H*_) to per capita ‘foraging’ rate, *f*(*L*_*H*_,*σ*) for three host genotypes. The candidate models for foraging rate, *f*(*L*_*H*_,*σ*), varied in complexity (Table 1). In model 1 (*null*), per capita foraging rate (*f*) is a single parameter (*f*). In model 2 (*size only*), a size-specific foraging coefficient (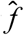) is multiplied by *L*_*H*_^2^, proportional to surface area, a common grazing model (Hall et al. 2007b; Kooijman 2010). In model 3 (*spores only, linear*) and model 4 (*spores only, power*), foraging rate drops as spores fill body volume, ∝ *L*_*H*_^3^ (i.e., as *σ*/*L*_*H*_^3^ increases, governed by coefficients α and γ). Models 3 and 4 both assume foraging does not scale with surface area. The most complex variants, model 5 (*size and spores, linear*) and model 6 (*size and spores, power*), combine surface area with the spore-mediated foraging depression. Models 1a–6a were fit assuming a shared foraging coefficient (*f* or 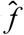) for both infected and uninfected classes. For models 1b–6b, we estimated parameters *f*_*j*_ or 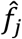 separately for infection class *j*. We inserted the best fitting function, assuming constant host length, into the dynamic epidemiological model (see *Dynamical Model* below; Figs. 2,A3). Additionally, the winning function enabled us to estimate depression of foraging during epidemics (see *Field Survey* below; Figs. 5,A4).

**Table 1.** Results of the model competition to estimate foraging rate, ***f***(***L***_***H***_, ***σ***). Models 1a–6a fit a common ‘foraging’ parameter (*f* or size-specific 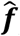 to infected and uninfected hosts together for each genotype. In models 1b–6b, foraging parameters (***f***_***j***_ or 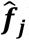) were estimated separately for uninfected (***f***_***S***_ or 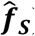) and infected (***f***_***I***_ or 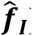) hosts in each genotype. Body length, ***L***_***H***_, and spore yield, *σ*, were measured empirically (Fig. 1d-f), and we estimated the linear sensitivity coefficient (*α*, mm^3^·spore^-1^) and power coefficient (*γ*) for each genotype.

| Model | Foraging<br>rate, $f(L_H, \sigma)^a$ | Pars<sup>b</sup> | AIC | $\Delta\text{AIC}^c$ | Akaike<br>weight ($w_i$)<sup>d</sup> |
| --- | --- | --- | --- | --- | --- |
| (6b) *Size and spores,*<br>*power* | $\hat{f}_j L_H^2 \left(1 - \alpha \left(\frac{\sigma}{L_H^3}\right)^\gamma\right)$ | 18 | -213.5 | 0.0 | 0.62 |
| (5b) *Size and spores,*<br>*linear* | $\hat{f}_j L_H^2 \left(1 - \alpha \left(\frac{\sigma}{L_H^3}\right)\right)$ | 15 | -212.5 | 0.9 | 0.38 |
| (3b) *Spores only, linear* | $f_j \left(1 - \alpha \left(\frac{\sigma}{L_H^3}\right)\right)$ | 15 | -96.3 | 117.2 | $2.2 \times 10^{-26}$ |
| (4b) *Spores only, power* | $f_j \left(1 - \alpha \left(\frac{\sigma}{L_H^3}\right)^\gamma\right)$ | 18 | -94.3 | 119.2 | $8.2 \times 10^{-27}$ |
| (2b) *Size only* | $\hat{f}_j L_H^2$ | 12 | -74.2 | 139.3 | $3.5 \times 10^{-31}$ |
| (6a) *Size and spores,*<br>*power* | $\hat{f} L_H^2 \left(1 - \alpha \left(\frac{\sigma}{L_H^3}\right)^\gamma\right)$ | 12 | -71.3 | 142.1 | $8.5 \times 10^{-32}$ |
| (5a) *Size and spores,*<br>*linear* | $\hat{f} L_H^2 \left(1 - \alpha \left(\frac{\sigma}{L_H^3}\right)\right)$ | 9 | -67.8 | 145.6 | $1.5 \times 10^{-32}$ |
| (1b) *Null* | $f_j$ | 12 | -6.1 | 207.4 | $5.7 \times 10^{-46}$ |
| (3a) *Spores only, linear* | $f \left( 1 - \alpha \left( \frac{\sigma}{L_H^3} \right) \right)$ | 9 | 136.5 | 350.0 | $6.2 \times 10^{-77}$ |
| (2a) *Size only* | $\hat{f} L_H^2$ | 6 | 143.6 | 357.1 | $1.8 \times 10^{-78}$ |
| (4a) *Spores only, power* | $f \left( 1 - \alpha \left( \frac{\sigma}{L_H^3} \right)^\gamma \right)$ | 12 | 144.7 | 358.2 | $1.0 \times 10^{-78}$ |
| (1a) *Null* | $f$ | 6 | 262.1 | 475.6 | $3.3 \times 10^{-104}$ |

<sup>a</sup> Per capita rates; technically, this is “clearance rate”. Units for $f_j$ : L·day<sup>-1</sup>; for size-specific $\hat{f}_j$ :
L·mm<sup>-2</sup>·day<sup>-1</sup>
<sup>b</sup> Number of parameters estimated for hosts of three genotypes, including a variance parameter estimated for each infection class (models 1b–6b) and genotype (all models). Parameters $\alpha$ and $\gamma$ were fixed at zero (i.e., not estimated) for uninfected hosts in models 3b–6b.
<sup>c</sup> The winning model has $\Delta\text{AIC} = 0$ . Models with $\Delta\text{AIC} > 10$ have essentially no support.
<sup>d</sup> The probability that the model is the best among those under consideration.

**Fig. 1.**
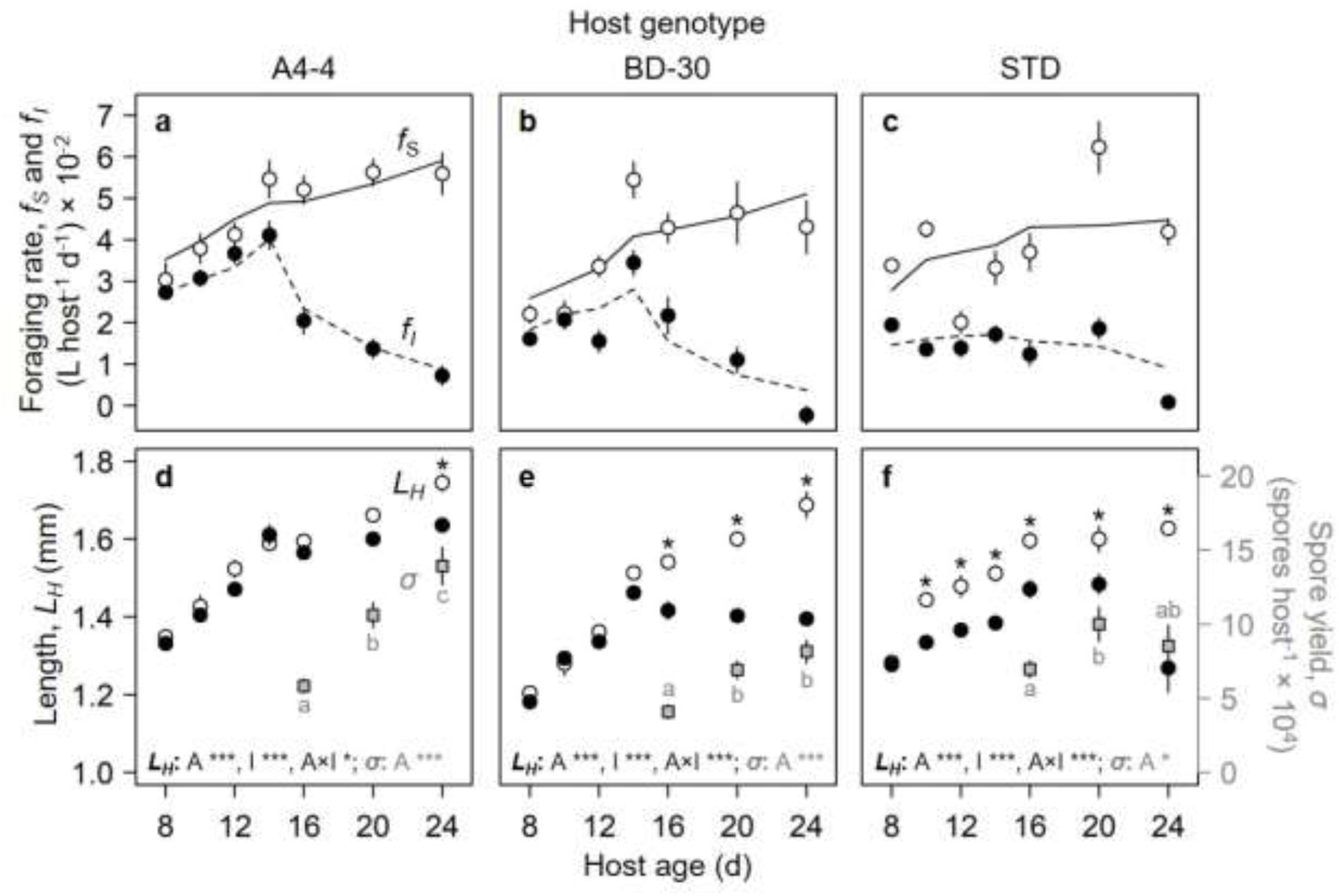
Parasites depress host foraging rate, f, as functions of host length, L_H_, and spore yield, σ. Foraging rate (a-c): Foraging rate (*f*, mean ± 1 SE) of uninfected (*f*_*S*_, white circles) and infected (*f*_*I*_, black circles; exposed to spores when six days old) individuals of three genotypes of a zooplankton host with the best-fitting foraging function (*f*_*S*_, equ. 2.a, solid lines; *f*_*I*_, equ. 2.b, dashed lines). (a,b) For genotypes A4-4 and BD-30, foraging rate increased with age (thus, body size) of uninfected hosts and those at early stages of infection. Foraging then dropped as infected hosts filled with spores. (c) Infection reduced foraging rate earlier for the STD genotype. *Spore yield and host length (d-f):* Host length (*LH*) of uninfected (white circles) and infected (black circles) hosts and spore yield (*σ*, grey squares) of three host genotypes: (d) A4-4, (e) BD-30, (f) STD. Spore yield also increased with age (noting a few [*N* = 3], smaller STD hosts at 24 days). *P*-values are from GLM-based tests of age (*A*), infection (*I*), and their interaction (*A* × *I*) on length, and of age on spore yield (p < 0.001 ***, p < 0.01 **, p < 0.05 *). Asterisks above body length points indicate significant post-hoc pairwise differences (Tukey’s) between infection classes. Letters denote significant post-hoc differences in spore yield between age classes. Points: means ± 1 SE.

**Fig. 2.**
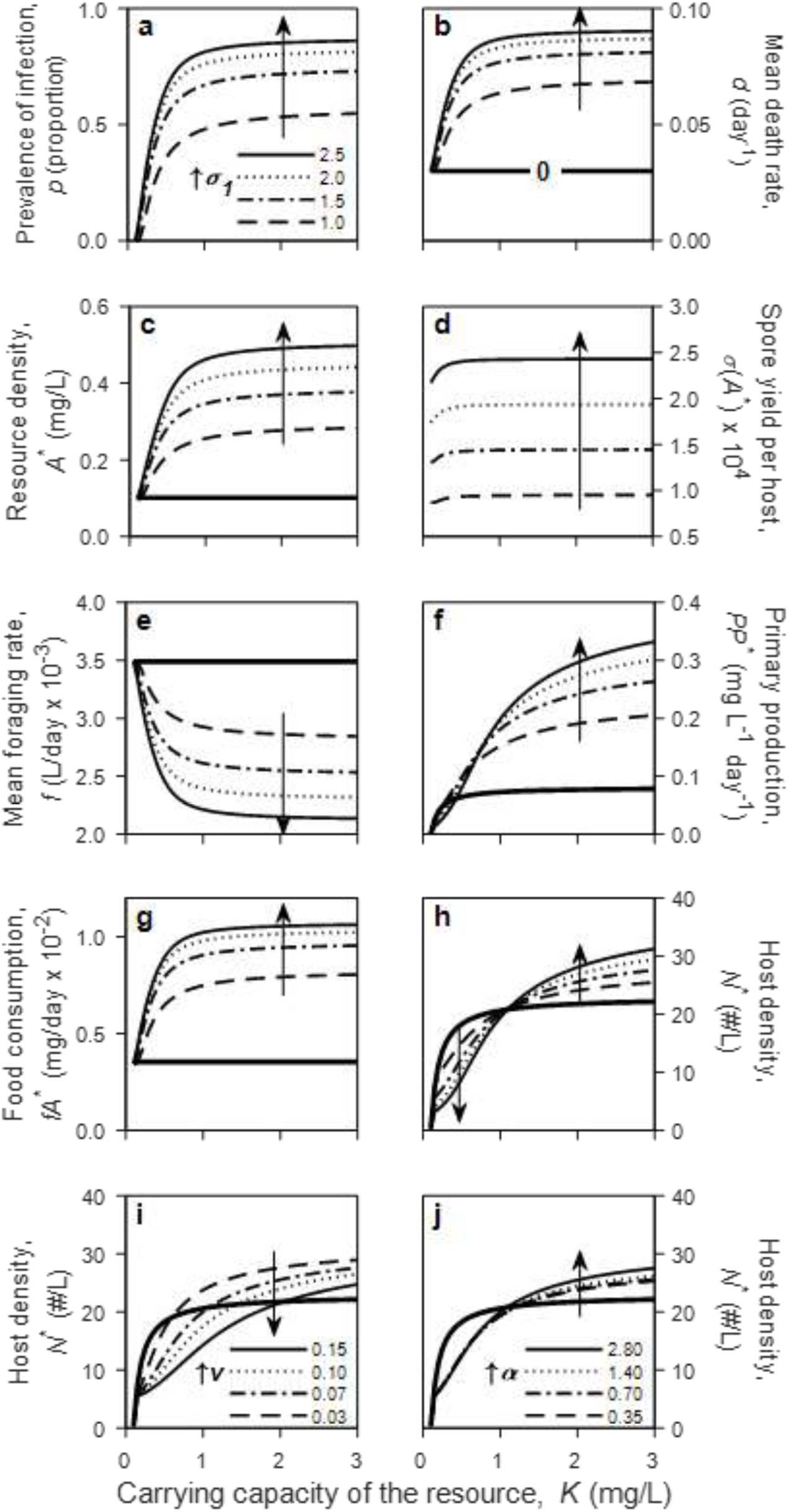
A fully dynamical model reveals a trait-mediated hydra effect through depression of foraging rate. Equilibrial density of hosts, *N*^*^, can increase during epidemics of a virulent parasite over gradients of carrying capacity, *K* (x-axis). Disease-free states are denoted by thick contours. For epidemics, arrows across contours show increasing values of maximal spore yield *σ*_1_ (spore·host^-1^·mg C^-1^×10^4^; panels a-f), virulence, *v* (day^-1^; panel *g)*, or sensitivity coefficient of foraging of infected hosts, *α* (mm^3^·spore^-1^; panel h). (a) Equilibrial disease prevalence (proportion infected, *p*^*\**^); (b) mean per capita death rate (*d+vp*^*^); (c) algal resources (*A*^*^); (d) spore yield (*σ*(*A*)); (e) mean foraging rate (*f* = [1-*p*^*^] *f*_*S*_ + *p*^*^*f*_*I*_; (f) primary production (*PP* = *r*_*m*_*A*^*^(1-*A*^*\**^/*K*); (g) resource consumption per host (*fA*^*^); and (h) total host density (*N*^*^). Hydras arise at higher *K* (*N*^*^ higher with disease [thin] than without) and become larger with higher *σ*_1_. The hydra effect was accentuated by (i) lower virulence on survivorship (*v* [day^-1^]) and (j) higher feeding sensitivity (*α* [×10^−5^ mm^3^·spore^-1^]). Therefore, hydras were more likely with higher carrying capacity of the resource (*K*) and for parasites that depress mortality less strongly (lower *v*) and foraging more strongly (higher *α*).

#### Parameterization and competition of the foraging function

We used maximum likelihood and information theoretic methods to parameterize and compete the foraging models, implemented with Matlab (version 7.8 R2009a; MathWorks). We estimated parameters by fitting a model of algal loss through time due to foraging (Sarnelle and Wilson 2008; Strauss et al. 2019):

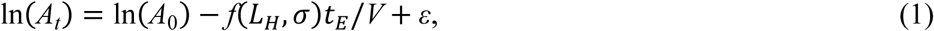

where *A*_*t*_ is the concentration of algae remaining at the end of the grazing period of length *t, A*_0_ is the concentration of algae in ungrazed reference tubes at time *t*_*E*_, *f*(*L*_*H*_,*σ*) is one of the foraging models (Table 1), *V* is the volume in the tube, and errors (*ε*) were normally distributed. (While it is technically ‘clearance rate’, the ‘foraging’ label avoids confusion with the immunological meaning of clearance.) We estimated parameters using maximum likelihood and competed models using standard information criteria (by calculating AIC, ΔAIC_*i*_, and Akaike weights, *w*_*i*_ for each model: see Table 1 (Burnham and Anderson 2002)).

The two best fitting models, 5b and 6b, fit the data equally well (ΔAIC < 1, Table 1, Fig. A1). We chose the more parsimonious model 5b (*size and spores, linear*) as the winner. We estimated 95% CI around the parameter of with 10,000 bootstraps (Table A1). We also compared parameters between host genotypes using 9,999 permutations (Gotelli and Ellison 2004) (Table A1). The slope and intercept of a regression of observed vs. predicted [ln(*A*_0_/*A*_*t*_)*V*/*t*] remaining algae were close to 1 and 0, respectively, indicating good performance (*Observed* = 1.007 × *Predicted* – 0.056 + ε; *R*^2^ = 0.55) (Piñeiro et al. 2008).

#### Outcome of the competition among foraging functions: results

Parasite infection reduced foraging rate of hosts, particularly during later stages of infection (Fig. 1a-c). In those later stages, fungal spores filled the body cavity of its host (Fig. 1d-f). Furthermore, infection stunted body size of sick hosts relative to uninfected hosts of the same age and genotype (Fig. 1d-f). As a result, the best fitting function for parasite-induced foraging depression (the ‘*size and spores, linear*’ model 5b; Tables 1,A1 and Figs. 1,A1) was:

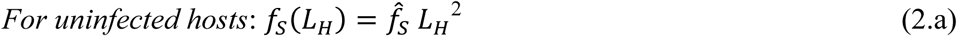

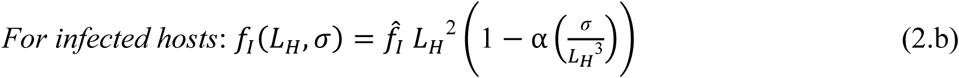

where 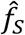 and 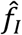 are ‘size-specific’ (size-independent) per capita foraging rates; *L*_*H*_ is host body length; *σ* is the spore load per infected host; and α is a linear sensitivity coefficient that governs depression of feeding as spores fill the host’s body cavity. These equations (equ. 2.a,b) capture how body size increased foraging. For uninfected hosts, foraging scaled with surface area (∝ *L*_*H*_^2^) (equ. 2.a; Fig. 1: solid lines, white circles). For infected hosts (equ. 2.b; Fig. 1: dashed lines, black circles), foraging rate also increased with surface area, though at a slower rate (since 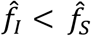; Table A1) but it decreased as their body volume (∝ *L*_*H*_^3^) filled with spores (*σ*) (Fig. 1).

### Dynamical Model: Foraging Depression Can Produce a Hydra During Epidemics

#### Structure of the dynamical model

We inserted the winning foraging function into a dynamical model. This model could then delineate conditions leading to foraging-mediated hydras vs. trophic cascades during epidemics (equ. 3, Table 2):

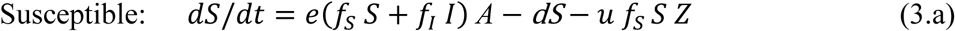

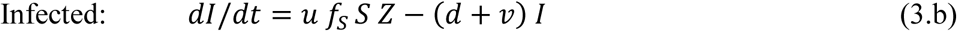

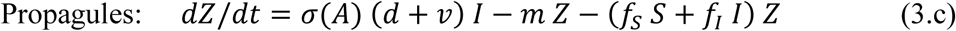

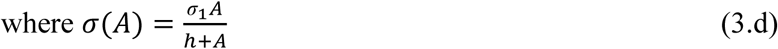

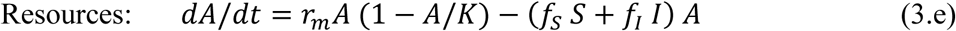

**Table 2.** Variables, parameters, and functions used in the dynamical host-parasite-resource model (equ. 3), with default values or ranges (when applicable).

| Symbol | Meaning | Default values/range | Units |
| --- | --- | --- | --- |
| $A$ | density of resource of host | --- | $\text{mg C} \cdot \text{L}^{-1}$ |
| $I$ | density of infected hosts | --- | $\text{host} \cdot \text{L}^{-1}$ |
| $N$ | density of hosts: $N = S + I$ | --- | $\text{host} \cdot \text{L}^{-1}$ |
| $S$ | density of susceptible hosts | --- | $\text{host} \cdot \text{L}^{-1}$ |
| $t$ | time | --- | day |
| $Z$ | density of parasite propagules (spores) | --- | $\text{spore} \cdot \text{L}^{-1}$ |
| $d$ | background death rate of hosts | 0.03 | $\text{day}^{-1}{}^a$ |
| $e$ | conversion efficiency of host | 8.5 | $\text{mg C}^{-1}{}^b$ |
| $L_H$ | body length of host | 1.4 | mm |
| $\hat{f}_S$ | size-specific foraging rate for susceptible<br>hosts | 0.0178 | $\text{L} \cdot \text{mm}^{-2} \cdot \text{day}^{-1}$ |
| $\hat{f}_I$ | size-specific foraging rate for infected<br>hosts | 0.0131 | $\text{L} \cdot \text{mm}^{-2} \cdot \text{day}^{-1}$ |
| $\alpha$ | sensitivity coefficient of foraging of<br>infected hosts | $2.86 \times 10^{-5}$ (Fig. 2h:<br>0.35-2.8 x $10^{-5}$) | $\text{mm}^3 \cdot \text{spore}^{-1}$ |
| $f$ | mean foraging rate of hosts: $f = (1-p)f_S +$<br>$pf_I$ | --- | $\text{L} \cdot \text{day}^{-1}$ |
| $f_S$ | foraging rate of susceptible hosts (equ. 2a) | 0.035 | $\text{L} \cdot \text{day}^{-1}$ |
| $f_I$ | foraging rate of infected hosts (equ. 2b) | 0 (bounded)-0.035 | $\text{L} \cdot \text{day}^{-1}$ |
| $h$ | half saturation constant, spore yield | 0.015 | mg C·L<sup>-1</sup> <sup>c</sup> |
| $K$ | carrying capacity of resources | 0.1-3.0 | mg C·L<sup>-1</sup> |
| $m$ | mortality of spores, $Z$ | 0.8 | day<sup>-1</sup> <sup>d</sup> |
| $p$ | prevalence of infection: $I/(S+I)$ | --- | unitless |
| $PP$ | primary production: $PP = r_m A(1-A/K)$ | --- | mg C·L<sup>-1</sup>·day<sup>-1</sup> |
| $r_m$ | max. per capita growth rate of $A$ | 0.8 | day<sup>-1</sup> <sup>a</sup> |
| $u$ | susceptibility | 0.0004 | host·spore<sup>-1</sup> <sup>e</sup> |
| $v$ | virulence on survival | 0.07 (Fig. 2g: 0.03-0.15) | day<sup>-1</sup> |
| $\sigma(A)$ | spore yield (equ. 3.d) | --- | spore·host<sup>-1</sup> |
| $\sigma_1$ | maximum spore yield | 1.0-2.5 x 10<sup>3</sup> (Fig.<br>2g,h: 1.5 x 10<sup>4</sup>) | spore·host<sup>-1</sup><br>·mg C<sup>-1</sup> |

<sup>a</sup> Reasonable value for this host and algae.
<sup>b</sup> Yields a reasonable instantaneous birth rate, $b$ , of 0.30 day<sup>-1</sup> for uninfected hosts at 1.0 mg C L<sup>-1</sup> (where $b = ef_S A$ ) (Hall et al. 2010).
<sup>c</sup> Reasonable value for this host (Strauss et al. 2015).
<sup>d</sup> A high loss rate due to solar radiation (Overholt et al. 2012) and other sources (e.g., consumption by non-focal hosts (Hall et al. 2009a)).
<sup>e</sup> Yields an infection risk (transmission rate, $\beta$ ) of 1.4 x 10<sup>-5</sup> L·spore<sup>-1</sup>·day<sup>-1</sup> (where $\beta = uf_S$ ).

Susceptible hosts (*S*, equ. 3.a) feed non-selectively at rate *f*_*S*_ on algal resources (*A*); infected hosts (*I*) feed at reduced rate *f*_*I*_. Feeding rates followed the best fitting foraging model described above (equ. 2.a,b). For simplicity, we assumed that hosts feed with a linear functional response. Ingested food is converted into offspring with efficiency *e*. Susceptible hosts (*S*) then die at background rate *d* or become infected following exposure (at rate *f*_*S*_) to spores (*Z*), with per spore susceptibility *u*. Infected hosts (*I*; equ. 3.b) die from infection (at enhanced rate *d* + *v*); they cannot recover. Spores (*Z*; equ. 3.c) are released from dead hosts; spore yield, *σ*(*A*), increases with algal resources (*A*) but saturates (with maximum *σ*_1_, and half-saturation constant *h*: equ. 3.d). Spores are lost at background rate *m* and via consumption by both host classes. Algae (equ. 3.e) grow logistically (at maximum per capita rate *r*_*m*_ and carrying capacity *K*) and are consumed by both host classes.

We simulated the model over a range of algal carrying capacity, *K*, and sensitivity of spore production to resources, *σ*_1_. We parameterized it using biologically reasonable values for this system (Table 2) and estimates of 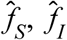 and *α* for the BD-30 genotype (equ. 2, Fig. 1, Table A1; assuming adult size *L*_*H*_ = 1.4 mm for both uninfected and infected hosts). Qualitatively similar results emerge using parameters from other genotypes. This dynamical model is not analytically tractable, thus we simulated it (using a standard adaptive step integrator in Matlab) for 1000 days. We then averaged densities of the state variables from *t* = 1000–2000 days. In the focal, biologically relevant region of parameter space shown here, the state variables reached a stable steady state by this time period. We found threshold combinations of *K* and mortality virulence (*v*) and of *K* and the sensitivity coefficient (*α*) that yielded foraging-mediated hydras. In each case, the threshold was found numerically (using a rootfinder) when host density at the boundary equilibrium (equ. A2) equaled host density at the interior equilibrium (*N*^*^ = *S*^*^ + *I*^*^, solved for numerically). Assuming lower baseline size-specific foraging of infected hosts 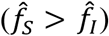, we found threshold levels of virulence mortality (*v*) at which hydra effects arose, either with sensitivity of foraging to spore accrual (*α* > 0) or not (*α* = 0). Then, at a given level of *v*, we found threshold levels of sensitivity to spore accrual (*α*) at which hydra effects arose, either due to both mechanisms of foraging depression 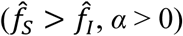 or only due to the effect of spores on 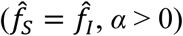.

#### Prediction of hydra from the dynamical model: results

Parasite-induced foraging depression can trigger a trait-mediated hydra effect (Fig. 2). Our model (equ. 3, Table 2) predicts that increasing the carrying capacity of algal resources (*K*, x-axis) or the maximum spore yield per infected host (*σ*_1_, contours) should increase equilibrial prevalence of infection (Fig. 2a). During larger epidemics, the average per capita death rate of hosts increases due to virulent effects of the parasite on host survivorship (Fig. 2b). Larger epidemics also yield greater density of resources, *A*, at equilibrium (*A*^*^, Fig. 2c). Since this density is also the minimal resource requirement of hosts, it increases with heightened mortality of hosts and foraging depression (Figs. A2,A3). More resources fuel greater within-host spore yield, *σ*(*A*) (equ. 3.d; Fig. 2d). Higher spore yield enhances spread of disease and boosts epidemic size, but it also depresses mean foraging rate of hosts, *f* (where *f* = (1-*p*^*^)*f*_*S* +_ *p*^*^*f*_*I*;_ *f*_*S*_ and *f*_*I*_ from equ. 2; Fig. 2e).

The model predicts either trophic cascades or foraging-mediated hydras – the outcome for host density depends on the relative effect of disease on resource production vs. on per capita resource consumption of hosts. The increase of resource density (due to virulent depression of foraging and survival) increases primary production (*PP*^*\**^ = *r*_*m*_*A*^*^(1-*A*^*^/*K*); see Appendix) – as long as *K* is high enough (*A*^*^ > *K*/2 – see Appendix Sections 2,3; Fig. 2f). Food consumption per host, *fA*^*^, also increases with *K* and *σ* (Fig. 2g). Host density, *N*^*^, then increases or decreases (relative to disease-free conditions) depending on the tension between responses of *PP*^*^ and *fA*^*^ (Fig. 2h; see also Appendix Sections 2,3; Fig. A3). At lower *K*, virulence on survival dominates, decreasing host density. At higher *K*, foraging depression and higher primary production increase host density. Therefore, the model predicts that larger epidemics may increase host density when parasites reduce feeding rate of their hosts enough in sufficiently enriched systems (see Appendix for more details). Furthermore, the foraging-mediated hydra effect should arise more readily when parasites are less lethal to their hosts (lower *v*; Figs. 2i, 3a,b), especially when infected hosts have lower baseline foraging rates 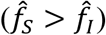 and their foraging is additionally depressed by within-host spore growth (α > 0, Fig. 3a; note 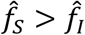 is enough to enable the hydra effect even when α = 0, Fig. 3b). Also, at a given virulence level (*v*), the hydra effect is more likely (i.e., can occur at lower *K*) when spore accrual more strongly suppresses foraging rate (higher *α*; Figs. 2j, 3c,d). The hydra effect occurs at lower α when 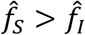 (solid line;) than when 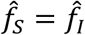 (dashed line; Figs. 3c,d) – therefore, both mechanisms of foraging depression (α > 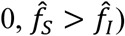 enhance the hydra effect. Finally, depression of host foraging rate may also drive higher infection prevalence (inferred from Fig. A3), through mechanisms involving higher spore production with higher resource density and lower per capita spore consumption (i.e., less removal of spores from the environment) by infected hosts (see Appendix and Fig. A3a-c.)

**Fig. 3.**
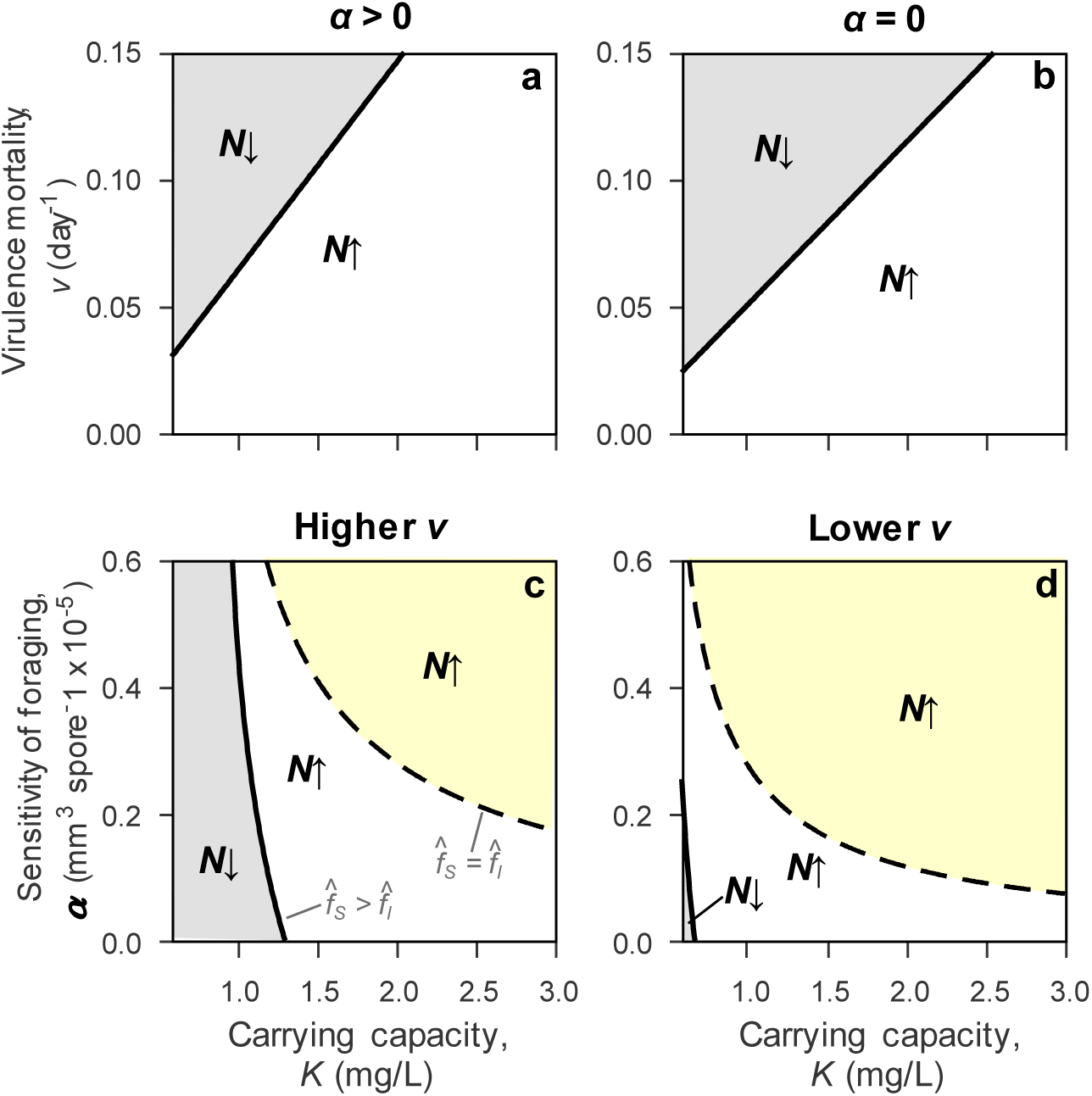
Parameter space predicting trophic cascades (host density decreases, N↓) or foraging-mediated hydra effects (N↑) over gradients of carrying capacity (K) of the resource. (a,b) Foraging-mediated hydras occur at a given *K* if virulence mortality, *v*, is not too high (below solid lines). Scenarios assuming susceptible hosts feed faster than infecteds 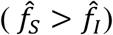: (a) foraging is sensitive to spores (α > 0) and (b) is not (α = 0). (c,d) Foraging-mediated hydras are predicted, at a given *K*, when the sensitivity coefficient, α, exceeds a threshold, which is smaller when susceptible hosts feed faster than infected hosts even without spore build up (i.e., 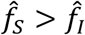, equ. 2; solid line, white and yellow region) than when they feed at the same rate without spores (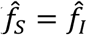, dashed line, yellow region alone). (c) Higher virulence on mortality (*v* = 0.07 day^-1^); (d) lower virulence (*v* = 0.03 day^-1^). All parameters follow defaults in Table 2.

### Field Survey: Evidence for the Hydra During Large Epidemics in Nature

#### Estimation of infection prevalence, algal density, spore yield: methods

We sampled 13 lakes in southern Indiana (Greene and Sullivan Counties, USA) weekly from August until the first week of December 2010. Here, we present data from the epidemic season (end of September through mid-November). On each sampling visit, we pooled three bottom-to-surface tows of a Wisconsin net (13 cm diameter, 153 µm mesh). From this sample, we estimated prevalence infection (*p*) by diagnosing at least 400 live *D. dentifera* at 20–50X magnification (Ebert 2005). From this sample, we estimated prevalence of infection in the adult size class only (*p*_*a*_). We also measured body length (*L*_*H*_) of uninfected and infected adult hosts (typically > 20 of each class). Additionally, we estimated the average spore yield (*σ*) of infected hosts (typically 5 to 40 hosts, pooled together). We estimated host density using preserved (60-75% ethanol) samples pooling three additional bottom-to-surface net tows. Finally, we indexed density of ‘edible’ (< 80 μm Nitex screening) algae in the epilimnion using narrow-band filters on a Trilogy fluorometer (Turner Designs) following chilled ethanol extraction (Webb et al. 1992; Welschmeyer 1994).

#### Index of ‘foraging depression’ and death rate: methods

For each lake population, we calculated an index of disease-induced ‘foraging depression’ of adult hosts using (1) prevalence and spore yield data (Fig. A4a), (2) body size of uninfected and infected adults (Fig. A4b), and (3) parameters from the winning foraging function (equ. 2; Table A1: 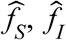, α; genotypes labeled 1–3 in Fig. A4b–d). We only summarize this calculation here (see Appendix Section 4 for details). For the infected class, we assumed that each infected adult shared the mean spore yield estimate for that lake-date (*σ*). With these parameters and data, we calculated mean foraging rate of adults, *f*_*a*,_ as mean foraging rate of each infection class of adults 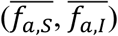 weighted by prevalence of infection of adults, *p*_*a*_; hence, 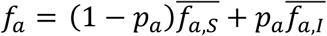 (see equ. A10.a-c). Next we calculated mean foraging rate of adults assuming differences in mean body size only, *f*_0_ (equ. A10.d). The index of foraging depression, *FD*, was then: *FD* = (*f*_*0*_ - *f*_*a*_) / *f*_*0*_ *100% (equ. A10.e; Fig. A4d). For each lake, we averaged this index, calculated at each sampling date, for each set of genotype-derived parameters (1–3); then, we averaged those three separate genotype-specific estimates to produce one value of *FD* per lake (see Fig. A4 for sample calculations). We also estimated average death rate of hosts during epidemics using the egg-ratio method (see Appendix Section 4 for details).

#### ‘Joint algal–host response’ index: methods

We calculated a ‘joint algal–host response’ index to test qualitative predictions of the dynamical model (i.e., hosts and resources should both increase during epidemics, particularly during larger ones). To quantify this index, we first estimated the linear slopes of hosts and algal resources through time (e.g., Fig. 4a,b). The ‘joint response’ index is the cross-product of these two vectors (Fig. 4c,d), estimated after standardizing their slopes (by the standard deviation in slope vectors among lakes). When both algae and hosts increased through time, the cross-product was positive (e.g., the large epidemic in Goodman Lake: Fig. 4a,c), consistent with a hydra effect. However, if only one of these (algae or hosts) increased through time, the cross-product was negative (e.g., the small epidemic in Long Lake: Fig. 4b,d). Densities of algae and hosts never both decreased through time (i.e., positive values only arose from two positive slopes, not two negative ones).

**Fig. 4.**
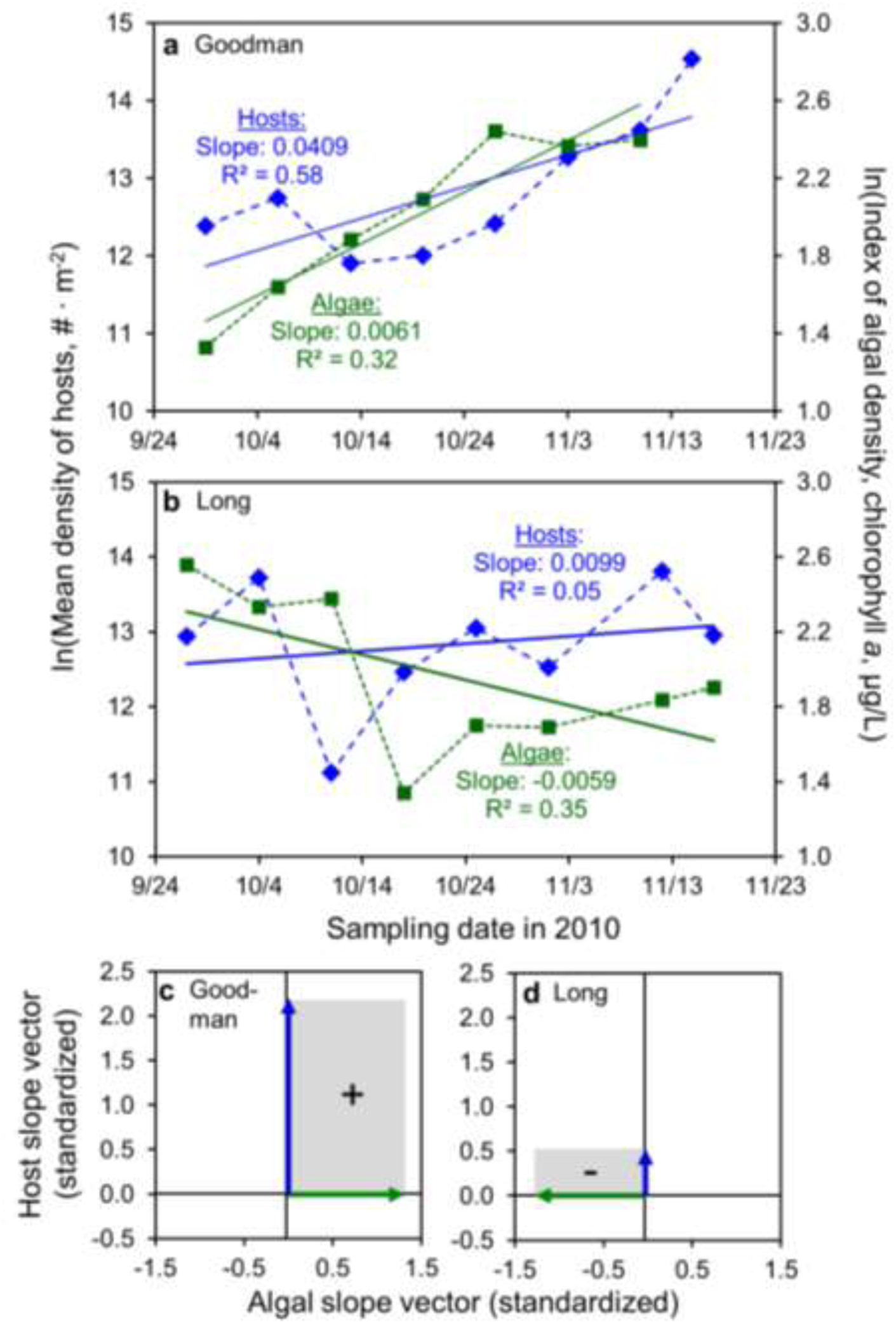
Changes in hosts and algal resources create a ‘joint algal–host response’. (a) During the large epidemic in Goodman Lake (max. infection prevalence: 48.6%; see also Fig. A4), both hosts and algal resources increased through time. (b) During Long Lake’s small epidemic (max. prevalence: 5.2%), hosts increased but algal resources declined. (c,d) The joint algal–host response index for (c) Goodman and (d) Long is calculated using cross products and the standardized temporal slopes (vectors). The ‘joint algal-host response’ index is the cross product of these vectors, i.e., their product (area), illustrated as grey rectangles. (This joint index is presented in Fig.5c,d.)

**Fig. 5.**
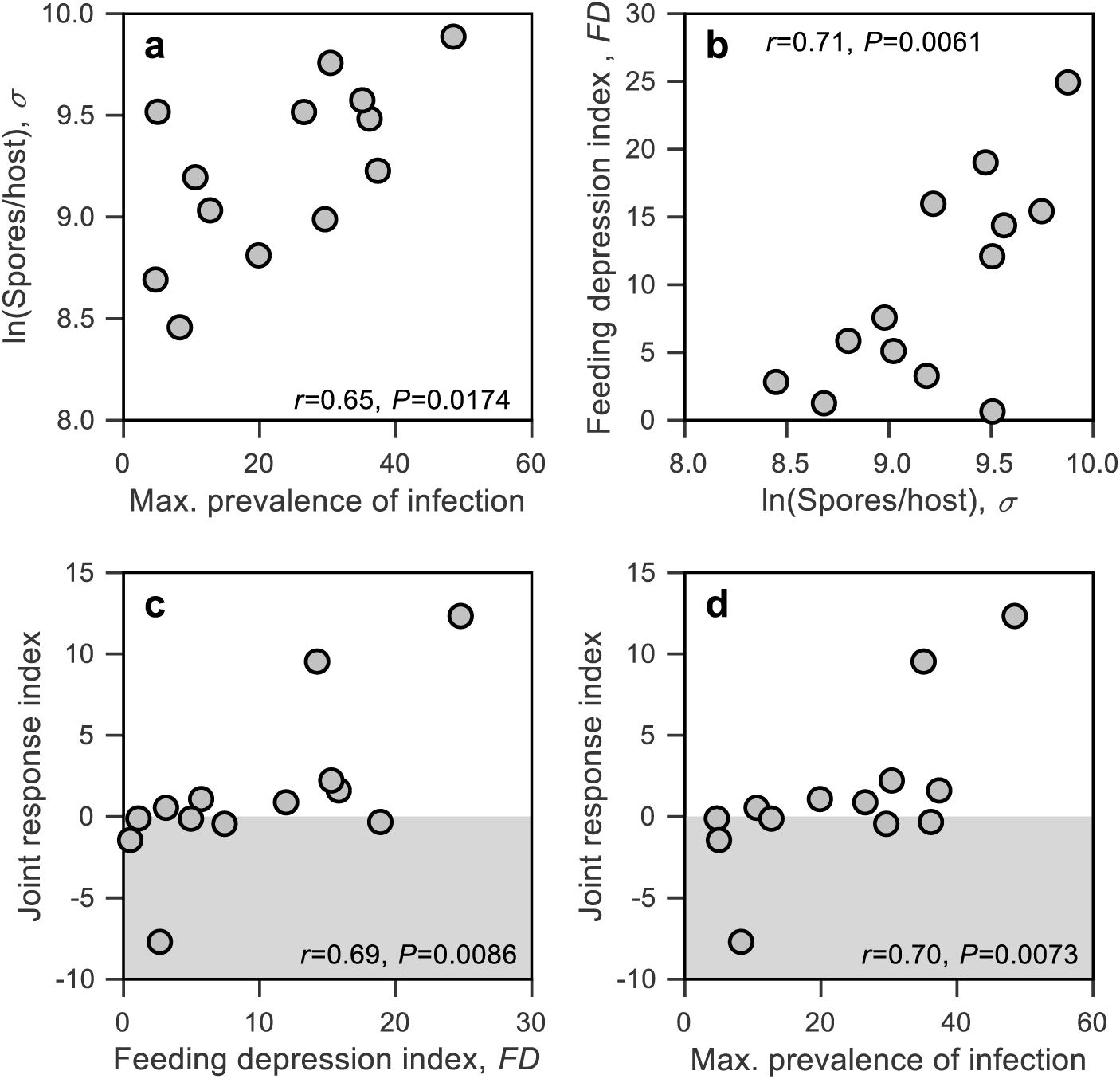
Evidence for a joint increase in densities of zooplankton (Daphnia) hosts and algal resources during natural fungal epidemics (a hydra). (a) Infected hosts produced more spores (*σ*), during larger epidemics (higher maximum prevalence of infection, *p*_max_). (b) These greater spore loads depressed average per capita feeding rate of adult hosts, *f* (calculated using length data; Fig. A4). (c) Stronger parasite-induced depression of foraging rate correlated with a larger index of joint algal–host response (Fig. 4c,d) through time during epidemics. (d) The joint algal–host response index was larger during bigger fungal outbreaks. Points are lake means. Pearson correlation coefficients (*r*) are accompanied by corresponding *P*-values.

#### Signature of the trait-mediated hydra effect in the field: results

In the field survey, we detected the hydra pattern anticipated by the dynamical model. Infected hosts yielded more spores in lakes with larger outbreaks (Fig. 5a) and more algal resources (shown previously (Civitello et al. 2015)). For lakes with greater spore loads, in turn, we estimated stronger depression of foraging by adult hosts (Figs. 5b,A4). Lakes with stronger foraging depression then had greater values of a dynamical index of algal resources and hosts through time (Figs. 4,5c). In this ‘joint algal–host response’ index, positive numbers indicate a hydra (see above). As predicted then, the signal of the foraging-mediated hydra effect arose during larger epidemics (higher prevalence) with stronger foraging depression (Fig. 5c,d). In contrast, mean death rate (estimate of *d* + v*p*) during epidemics was not correlated with the index of foraging depression (*R* = 0.23, *P* = 0.42) or the joint algal-host index (*R* = 0.26, *P* = 0.35).

## Discussion

Undeniably, large epidemics of virulent parasites can depress host densities. However, here we show that indirect feedbacks between hosts and resources can drive the opposite pattern: increased host density during outbreaks. More specifically, parasites that virulently depress infected host foraging rate can indirectly produce more hosts under certain conditions. Using a zooplankton-fungus-algal system, we show how infection by a virulent parasite depresses foraging of infected hosts. Then, using a dynamical model of host-parasite-resource interactions, we show when the foraging-mediate hydra effect should and should not arise. The model predicted hydras during larger epidemics that strongly depress foraging of hosts while, at the same time, not depressing fitness too much. We then turned to naturally occurring epidemics, finding support for a foraging-mediated hydra effect. During larger epidemics, more spores accumulated in host bodies, which depressed foraging. Reduced foraging, in turn, correlated with a joint increase in hosts and algal resources – a signature of the hydra effect.

How and why does the foraging depression mechanism work? In the model, it works via two components of host density: the ratio of resource production and resource consumption per host. Both components start with an increase in the minimal resource requirement of hosts (an indirect effect). Hosts require enough resources to offset increased mortality (resulting from parasite virulence) with reproduction (extending logic from (Grover 1997)). Reduced foraging further increased this requirement. The subsequent increase of resource density can increase resource production (Case 2000). However, higher food density compensates for slower feeding, yielding no net change in resource consumption. Therefore, foraging depression alone enhances the likelihood of hydra effects during epidemics. In this system, foraging depression arose in multiple ways. Infected hosts had lower size-specific feeding rate; infected hosts were smaller (reducing size-dependent feeding further); and spore accumulation in host bodies substantially diminished foraging. Higher density of resources should exacerbate this spore-accumulation effect (Civitello et al. 2015; Hall et al. 2009b). Finally, these hosts slow feeding when contacting parasite propagules (Hite et al. 2017; Strauss et al. 2019). Hence, in this plankton system, multiple mechanisms produce foraging depression. Since parasite-mediated foraging depression arises commonly in other systems as well (Hite and Cressler 2019; Hite et al. 2020), this trait-based mechanism for a hydra may apply quite broadly.

Even with foraging depression, hydra effects may still not arise unless additional conditions are met. First, hosts must strongly control their resource. While *Daphnia* famously depresses its resources, not all hosts can (Borer et al. 2005; Shurin and Seabloom 2005). Second, the subsequent increase in resource density must enhance resource productivity. Some resources follow a more donor-controlled, chemostat-style supply (Polis et al. 1997); in these cases, productivity drops as resource density increases, eliminating the hydra (see Appendix). Notably, many experiments impose donor control, which means foraging-mediated hydras cannot occur. Furthermore, sufficient enrichment is needed for higher density to increase resource productivity. Third, parasites cannot depress survivorship or fecundity (Hall et al. 2007a; Lafferty and Kuris 2009) too strongly. Those forms of virulence increase per host resource consumption, potentially overwhelming any increase in resource productivity. Fourth, epidemics must become large enough to trigger the requisite indirect effects to densities and traits. While this is a lengthy set of requirements, our results suggest they happen in at least this planktonic system. It remains to be determined how many other systems can also produce a foraging-mediated hydra.

Where does this foraging-mediated hydra result fit within other behaviors of host-parasite-resource systems? First, hydras can arise via other mechanisms (Abrams 2009; Abrams and Matsuda 2005; Cortez and Abrams 2016). Increased mortality of hosts during epidemics could stabilize oscillatory host-resource cycles to increase host density. Here, the linear functional response yielded stable dynamics, obviating evaluation of this mechanism. Yet, we found no relationship between mean per capita death rate and the joint algal-host index (but see (McIntire and Juliano 2018) for an example of increased mortality driving higher density in mosquitoes). Second, parasites can drive trophic cascades (Buck and Ripple 2017). In our model, cascades were more likely at lower productivity, for less sensitive foragers, and for more virulent parasites; trophic cascades have been found in our plankton system, too (Duffy 2007). Third, parasites can trigger ‘biomass-overcompensation’ in their host. This outcome, assuming certain trait asymmetries between life stages of hosts, can increase biomass of the life stage most readily infected (de Roos and Persson 2013; Preston and Sauer 2020; Schröder et al. 2009). Hopefully, a coherent theory will emerge that synthesizes these possibilities for hydras, cascades, and biomass overcompensation during epidemics.

Moving one step further, the foraging-mediated hydra effect should be integrated into a broader theory for the community ecology of disease. First, foraging depression by parasites should stabilize host-resource oscillations, providing another mechanism to produce a hydra effect (e.g., (Hilker et al. 2009; Hurtado et al. 2014)). Second, other food web interactors might stifle this foraging-mediated hydra. For instance, competitors of hosts could fix resources at their own minimal resource requirement (analogous to systems with inedible producers: (Grover 1995)). Therefore, competition might prevent hydras. Third, hosts can evolve during epidemics (Boots et al. 2009; Duffy and Forde 2009). This *Daphnia* host shows foraging-mediated relationships between fecundity and transmission rate (Auld et al. 2013; Hall et al. 2010) and between feeding rate and sensitivity to contact with spores (Strauss et al. 2019). Such relationships could interact interestingly with foraging-mediated hydras as hosts evolve during epidemics. Therefore, integration of the foraging-mediated hydra effect awaits future developments.

The foraging-mediated hydra effect means that large outbreaks may not depress host density. Parasite-mediated foraging depression occurs in a diverse array of systems (Hite and Cressler 2019; Hite et al. 2020). Yet, the foraging-mediated hydra here rests on a number of requirements, including that hosts strongly control resources, that resource productivity increases, and that infection only moderately increases mortality. It remains unknown how many other systems meet these conditions. However, it is important to note that these foraging-mediated hydra effects may produce desirable or undesirable outcomes. Hydras might prevent worrisome collapses in host density during large outbreaks. Yet, they also increase density of infected hosts, potentially elevating disease risk to humans (via contact with infected hosts) or spillover to other hosts. Future efforts should evaluate the frequency and magnitude of foraging-mediated hydra effects and their influence on disease and communities.

## Acknowledgements

We thank J.J. Potter for lab assistance. K. Boatman, Z. Brown, D. Grippi, J. Hite, C. Searle, and A. Smith helped in the field and lab. C. Gowler, J. Marino, C. Shaw, and C. Wood provided comments on earlier drafts of the manuscript. This project was supported by NSF (DEB-0841679 to MAD and DEB-0841817 and 1120316 to SRH, and Graduate Research Fellowships to RMP and MSS). We appreciate the cooperation of S. Siscoe (Division of Forestry) and R. Ronk (Division of Fish and Wildlife) at the Indiana Department of Natural Resources for access to field sites.

### Online Appendix

#### Overview

In the first section of this Appendix, we provide more results from the foraging assay (Fig. 1a-c). The spore yield and length data (Fig. 1d-f) were used to parameterize the various competing functions of foraging (as visualized in Fig. A1). All of the details of the winning model (Table A1) and of the competition itself (Table 1) are also shown here. Second, we provide an in-depth analysis of the response of host density to depression of foraging rate in the absence of disease (Fig. A2). Since it analytically conveys key logic for the more complex model in the main text, we describe it in some depth. Third, we study how parasite-induced foraging depression affects equilibrial densities of resources and hosts during epidemics, using key comparisons (Fig. A3). We contrast the dynamical model from the main text (equ. 3; Fig. 2), where parasites reduce both host survival and foraging, with two ‘virulence variants’ (described below). Finally, we describe methods and illustrate calculations used to describe foraging depression and mortality during epidemics (Fig. A4).

#### (1) More methods and results from the foraging rate experiment, and the parameterization and competition of foraging functions

##### Additional methods

We measured foraging rate, body size (host length, *L*_*H*_), and spore density per host (*σ*) to parameterize the foraging models for uninfected and infected hosts of three genotypes. Hosts and parasites were originally from lakes in Barry County, MI, USA, except one host genotype, Beaver Dam 30 (BD-30), which was from Greene County, IN, USA. To standardize maternal effects, each genotype was reared in Artificial *Daphnia* Medium (ADaM (Klüttgen et al. 1994)) mixed with filtered water from Lake Lanier (Georgia, USA), and fed 0.9 µg C mL^-1^ day^-1^ (a standard, non-limiting level) of a nutritious green alga (*Ankistrodesmus falcatus*). We generated cohorts of 8-, 10-, 12-, 14-, 16-, 20-, and 24-day-old animals by collecting them within a 24-h period (grouped as 10 per 150-mL beaker, kept at 20 °C in a 16:8 h light:dark cycle, then later spread to six per beaker at six days old). Exposed beakers then received parasites (900 spores mL^-1^). We transferred all hosts to fresh medium after the 24-h exposure, and then every 4 days until the day of the foraging rate assay. For the assay, hosts were placed singly into 15-mL centrifuge tubes. For each treatment, sample size exceeded 12 for each age × infection × genotype combination except *n* = 3 for 24-day-old infected STD hosts (most of these hosts had already died of infection by then). Hosts grazed on 0.45 µg C mL^-1^ of *A. falcatus* for 4 h; tubes were inverted every 20 min to ensure algae stayed suspended. Hosts were then removed from each tube and measured from the middle of the eye to the base of the tail spine at 40X. We quantified food remaining in the tube using a Trilogy fluorometer (*in vivo* module, Turner Designs, Sunnyvale, CA, USA).

To estimate foraging rate of the infected class, we only used hosts that developed infections that reached the ascospore stage (Stewart Merrill and Cáceres 2018). Spores were visually apparent once infected hosts were 16+ days old (i.e., 10+ days post-exposure). To estimate spore yield, we transferred hosts to microcentrifuge tubes, gently smashed each individual using a pestle, and counted the released spores using a hemocytometer at 200X magnification. Since infected hosts less than 16 days old typically contain very few spores (Auld et al. 2014), we assumed they contained none during the assay, but diagnosed them later, retaining only successfully infected hosts in the analysis. After removing hosts that did not ultimately develop infections, each treatment had at least 9 replicates, except the 24-day-old infected STDs (discussed above).

##### Additional results

Individuals from the three host genotypes grew through time when uninfected, but growth tended to slow or plateau once spores accumulated in their bodies (i.e., 10+ days post-exposure to the parasite, when hosts were 16+ days old; Fig. 1d-f). The best fitting models (5b: *size and spores, linear*, and 6b: *size and spores, power*) explained the drop in foraging of infected hosts (Fig. 1,A1; Table 1). Parameter estimates for the winning model varied among the three genotypes (Table A1), and that variation was included in calculations of the index of foraging depression in the lakes (Fig. 5b,c; Fig. A4c,d).

#### (2) Theoretical insights from the disease-free subsystem of the dynamical model

The disease-free subsystem only has hosts (all of whom are uninfected, or susceptible, *S*) and resources (*A*). Imagine that the host feeds with a linear functional response (at foraging rate *f*; following the model in the main text) while the algal resource grows logistically, yielding:

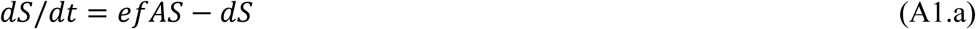

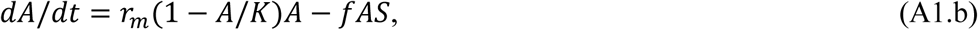

where population growth rate of susceptible hosts (*dS*/*dt*, equ. A1.a) increases with foraging rate *f* and conversion of consumed resources into offspring (with efficiency *e*) but decreases at a background loss (death) rate *d*. Per capita birth rate, *b*, is *e f A* and equals death rate *d* at equilibrium. The growth rate of the resource (*dA*/*dt*, equ. A1.b) is logistic as governed by maximal growth rate *r*_*m*_ and carrying capacity *K*. Without grazing, the algal resource would reach its carrying capacity (*K*) at the boundary equilibrium; with grazing, the interior equilibrium is:

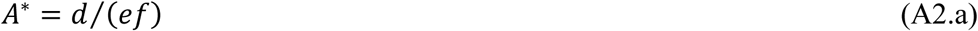

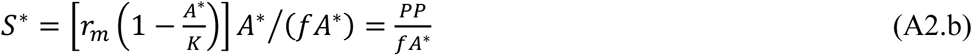

At this equilibrium, resource density (*A*^*^, equ. A2.a) is equal to the minimal resource requirement of the host (where low death rate, *d*, high conversion efficiency, *e*, and/or high foraging rate, *f*, lead to strong control over the resource, or lower *A*^*^). Here, per capita death rate (*d*) of the host equals its per capita birth rate, *b* (where again, *b* = *e f A*^*^). Equilibrial host density (*S*^*^, equ. A2.b) is the ratio of two key quantities (written in a particular way here to maximize meaning below). The numerator of this ratio is primary production of the algal resources, *PP* = *r*(*A*^*\**^) *A*^*\**^; that is, per capita productivity of the resource, *r*(*A*^*\**^) = *r*_*m*_ (1 - *A*^*^ / *K*) (in square brackets of equ. A2.b) times equilibrial algal density, *A*^*^. Primary productivity, *r*(*A*^*\**^) *A*^*\**^, follows the familiar, unimodal hump of the logistic model with increasing *A*; thus it is maximized at *K*/2. The denominator is per host consumption of the resource, *f A*^*^ (which itself is proportional to host per capita birth rate, *b*). Host density, *S*^*^, then depends on how primary productivity (PP, the numerator) is partitioned among grazers (the denominator, with each grazer taking portion *f A*^*^).

How will this interior equilibrium (equ. A2) respond to depressed foraging rate, *f*? We can see that a slower forager (‘lower *f*’) can reach higher density than a faster forager (‘higher *f*’) if carrying capacity is above a certain threshold (this threshold level of *K* increases with death rate [Fig. A2a,b]). More insight arrives from a tiny bit of calculus. Not surprisingly, resource density will *always* increase if *f* drops since:

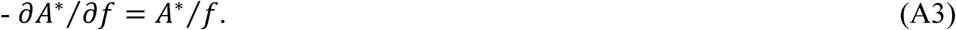

(Notice the negative partial derivative here to denote foraging rate, *f*, shrinking.) Hence, a slower foraging host needs a higher minimal resource requirement. However, a drop in foraging rate (*f*) can either elevate or depress density of the host, *S*^*^. The outcome depends on how primary productivity, *r*(*A*^*^) *A*^*^, responds to the increase in resource density, *A*^*^ (given lowered foraging rate; equ. A3) and how each host’s rate of resource consumption, *f A*^*^, changes. This latter rate does not change with decrease in *f* alone (i.e., -∂(*fA*^*^)/∂*f* = 0) because the depression of foraging rate is exactly offset (compensated) by an increase in resource density (given equ. A3). Thus, the response of host density to foraging depression solely hinges on how primary productivity changes with increased crowding of resources. Formally,

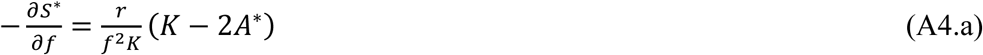

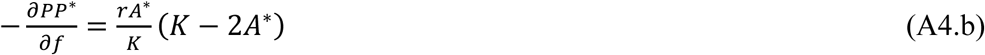

which says that hosts, *S*^*^, will increase with a drop in foraging, *f* (i.e., -∂*S*^*^/∂*f* > 0) if resource density (*A*^*^) is less than half of the resource’s carrying capacity (*K*/2; equ. A4.a). That threshold happens when *A*^*^ is below peak primary productivity (*K*/2) – notice how primary production also increases with a drop in foraging in this same range (equ. A4.b). Therefore, higher primary productivity can support higher density of the more slowly foraging hosts (yellow region, ‘*S*↑’; higher *f-K* space in Figs. A2c,d; more *f-d* space at higher *K* in Figs. A2e vs. A2f). This case is most likely when the host strongly controls the resource (i.e., when its minimal resource requirement, *A*^*^, is well below *K* due to high initial foraging rate or in more productive systems (higher *K*; Figs. A2c,d). Alternatively, when *A*^*^ > *K*/2, total primary productivity drops as resources increase due to foraging depression. Lower primary productivity supports fewer hosts. This case arises when hosts do not control their resources strongly (i.e., then they cannot depress *A*^*^ below *K*/2 due to low foraging rate or in less productive systems (white regions, ‘*S*↓’; lower *f-K* space in Figs. A2c,d; more *f-d* space at lower *K* in Figs. A2e vs. A2f). Note also that a resource that is donor-controlled / has chemostat-style renewal would prohibit an increase of hosts with declining feeding rate. For instance, imagine that resource renewal was *a* (*A*_*S*_ – *A*) in equation A1.b (with dilution rate *a* [day^-1^] and supply point *A*_*S*_ [mg/L]) instead of *r*_*m*_ *A*(1-*A*/*K*). In this case, the minimal resource requirement would not change (*A*^*^ = *d*/(*ef*)), and primary production (*PP* = *a* [*A*_*S*_ – *A*^*^]) would always decline with decreasing f (-∂PP/∂f = - *a A*^*^/*f*) as would host density [*C*^*^ = (*a*/*f*)(*A*_*S*_/*A*\* - 1); -∂*C*^*^/∂f = -*a*/*f*^2^].

This equilibrial analysis of the susceptible host/grazer (*S*)–algal resource (*A*) subsystem prompts the following qualitative predictions for a more complicated system with a parasite that depresses foraging rate of its host (equ. 2-3).

1. Algal density, *A*^*^, should always increase when foraging rate of the host, *f*, drops.
2. However, the response of host density depends upon how primary productivity, *r*(*A*^*^) *A*^*^, responds to this higher density of resources (i.e., whether *A*^*^ is higher or lower than peak primary productivity, *A*^*^ = *K*/2).
3. Primary productivity should increase, and hence host density should increase, with depressed foraging rate when hosts strongly control their resource (relatively low *A*^*^) or in more productive systems (higher *K*; case 1 in Fig. A2b). Here, we see a joint increase (*A*^*^↑, *S*^*^↑) caused by foraging depression.
4. Primary productivity declines and host density drops with foraging depression when grazers cannot strongly control their resource (relatively high *A*^*^, due to high *d*, low *e*, and/or low baseline *f*; case 3 in Fig. A2a) or in less productive systems (lower *K*; case 3 in Fig. A2b). Then, we expect a trophic cascade-like pattern (*A*^*^↑, *S*^*^↓).
5. However, per capita resource consumption, *f A*\*, and hence per capita birth rate (*b* = *e f A*^*^), should not change with a decline in foraging rate, *f*, alone. In other words, the response of host density to depressed foraging does not involve per capita birth rate. This aspect is critical, since host response depends on how decreasing *f* affects primary production (predictions 3 and 4) as well as per capita resource consumption (equ. A2.b).
6. This dependence on per capita resource consumption explains why host density should decline with increased death rate, *d*, i.e., if the parasite virulently depresses survival. If death rate increases:

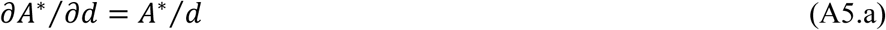

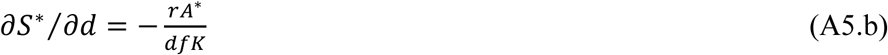

thus, resource density should increase (equ. A5.a) but grazer density should decrease (equ. A5.b) at higher *d*. The response of producer density to higher *d* qualitatively echoes that seen for foraging depression (equ. A3). However, a little bit of calculus shows that host density always declines with increasing *d* – even if primary production increases (i.e., *A*^*^ < *K*/2) – because per capita consumption by grazers has to increase too much with higher *d* (since *e f A*^*^ = *b* = *d* at equilibrium, by definition). In other words, with increasing *d*, higher consumption demands per host overwhelm any primary productivity response. This insight explains why the same *f-K* combination at low *d* produces more hosts with lower *f* but fewer hosts at higher *d* (contrast dot in Fig. A2c [lower *d*] vs. A2d [higher *d*]). Similarly, at low *K*, an *f-d* combination that would produce less hosts with foraging depression yields more hosts at higher *K* (contrast dot in Fig. A2e [lower *K*] vs. A2d [higher *K*])
7. Hence parasite-mediated hydra effects become more likely at higher productivity for hosts which control their resources (guaranteeing *A*^*^ < *K*/2). Furthermore, they are more likely when foraging depression is large (boosting *PP*) but when parasites are not too virulent (lower *v*, which would increase food consumption, *fA*^*^, per host too much, overwhelming gains in PP).

#### (3) Further discussion of the dynamical model of the full host–parasite–resource system

In addition to the modeling results presented in the main text, we examined two other ‘virulence variants’ to better understand the predicted response of hosts to either foraging depression alone (lower *f*) or higher virulence on survival (*v*) alone (Fig. A3). These examples build on the intuition from the disease-free subsystem (section 2). The first variant (left column, Fig. A3) extends the example in the main text: during epidemics, hosts experience foraging depression (sensitivity coefficient α > 0; see equ. 2.b) and virulence on survival (*v* > 0). ‘Variant 2’ features the same virulence on survival but no foraging depression (α = 0, *v* > 0; middle column). ‘Variant 3’ models only foraging depression without virulence on survival (α > 0, *v* = 0; right column).

These three ‘virulence variants’ disentangle the effects of decreased foraging and survival on epidemiology (i.e., disease prevalence at equilibrium) as well as densities of resources and hosts. **Disease prevalence** (proportion infected, *p*^*\**^) at equilibrium is quantitatively different among the variants (Fig. A3a–c). At a given carrying capacity (*K*) and maximal spore yield (*σ*_1_), prevalence is typically greater in ‘variant 1’ and ‘variant 3’ (which include foraging depression), compared to ‘variant 2’ (which only includes virulence on survival). Therefore, all else equal, parasite-driven foraging depression promotes larger epidemics through a combination of greater total host density (contrast Fig. A3m,o vs. A3n) and less removal of spores from the environment by already-infected hosts. Disease prevalence in ‘variant 1’ is also enhanced by greater resource density (contrast Fig. A3d vs. A3e) – and thus spore yield of infected hosts – relative to ‘variant 2.’ The **resource response** is striking but unidirectional. Resource density, *A*^*^, is:

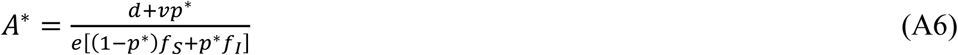

which is the ratio of per capita mortality, *d* + *vp*^*^ to per capita, per resource birth rate, i.e., conversion rate, *e*, times mean feeding rate of the population, taking into account proportion susceptible, (1-*p*^*^)*f*_*S*_, and infected, *p*^*^*f*_*I*_ (equ. A6). In ‘variant 1’ (Fig. A3d), *A*^*^ shows synergy between the indirect effects of virulence on survival (Fig. A3e) and foraging depression (Fig. A3f) on resource density. The effects of foraging depression alone on *A*^*^ are actually small (given the parameters). **Primary production**, *PP*, largely mirrors the algal density response (Fig. A3g–l). As described in the second section of this supplement (*Theoretical insights from the disease-free subsystem* above), primary production is *r*_*m*_ *A*^*^(1 - *A*^*^/*K*), and it determines the numerator of host density. Over much of the *K* range, primary production increases during epidemics (more subtly with only foraging depression [Fig. A3j], more for virulence on survival [Fig. A3h], and synergistically for both [Fig. A3g]). **Food consumption**, *fA*^*^ = (1-*p*^*^)*f*_*S*_ + *p*^*^*f*_*I*_, follows a relatively similar pattern. It is highest when both α>0 and *v*>0 (Fig. A3j), lower when infection only imposes mortality (Fig. A3k), but does not change when it only imposes foraging depression (because foraging is compensated for in the minimal requirement exactly; Fig. A3l). **Total host density**, *N*^*^, is:

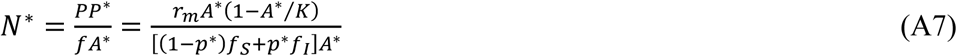

the ratio primary production to food consumption (equ. A7). It reflects tension between the two sources of virulence. With only virulent effects on survival, host density decreases (Fig. A3n). In contrast, over the vast majority of the *K* gradient (except for very low *K*), foraging depression alone indirectly increases host density during epidemics (Fig. A3o), given that these hosts strongly control their algal resource (i.e., they have low minimal resource requirement, *A*^*\**^). Thus, the response of host density to a combination of foraging depression and virulence on survival depends on carrying capacity. At lower *K*, the host-decreasing effect of virulence on survival dominates; at higher *K*, the host-increasing effect of foraging depression prevails (Fig. A3m). This result reminds us that the response of host density during epidemics does not merely follow an increase in primary production. The per capita foraging consumption (*fA*^*^) by hosts needed to ‘break even’ determines how many hosts the primary production can support. Hosts suffering higher virulence on survival require higher resource consumption to break even; hosts experiencing foraging depression do not (using the logic in section 2 above).

In summary, this comparison of model variants clarifies the response of hosts and their resources during epidemics. If the parasite only virulently lowers survival of hosts, the model predicts only a trophic cascade, where resources increase while host density declines relative to disease-free conditions. (If the consumer–resource system could oscillate, host density might increase with lower survival, under some conditions (Abrams 2009). This possibility was not modeled here.) In contrast, if the parasite only lowers foraging rate, a host with these *Daphnia*- like traits (i.e., exerting strong control over its resource; Table 2) should typically increase during epidemics. In other words, both host and resources increase, relative to disease-free systems. Parasites that depress host feeding rate should also typically have larger epidemics compared to parasites that reduce survival only. For parasites that both depress survival and foraging rate, the host response depends on the relative strength of effects of survival vs. foraging rate on host density. This balance can shift with carrying capacity of the resource, as increases in host density during epidemics are more likely when carrying capacity is higher.

#### (4) Field survey: more methods and sample calculations of indices describing data from the field survey

##### Methods for calculating death rate

In the field survey, we calculated temperature-dependent death rate in a way that incorporates diel migration of the host. This species of host typically migrates below the thermocline (into the ‘metalimnion’) of lakes during day into deeper, colder, but still oxygenated (> 1.0 mg/L dissolved O_2_ [DO]) waters. Then, at night, it moves above the thermocline into upper, warmer habitat (the ‘epilimnion’) (e.g., (Duffy et al. 2005; Hall et al. 2005). Therefore, using temperature data, we calculated depth of the thermocline (during periods of stratification) by: (1) converting temperature data into densities (following (Chen and Millero 1977)); (2) then calculating buoyancy frequency, *N* = (*g*/*ρ*(*dρ* /*dz*))1 2 [where *g* is acceleration due to gravity, *ρ* is mean density, and *dρ/ dz* is the vertical density gradient], at 0.1 m depths by differentiating piece-wise cubic splines fit through the density-depth data (with pchip.m in Matlab); and (3) finding the thermocline as the depth of maximum buoyancy frequency. We found the oxygenation threshold (1.0 mg DO) using cubic splines fit through DO-depth data. With temperature, thermocline depth, and oxygenation threshold information, we calculated mean development time in the oxygenated metalimnion (day, *D*_*M*_) and epilimnion (night, *D*_*E*_).

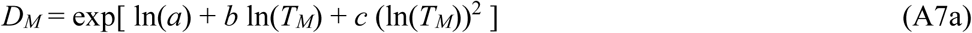

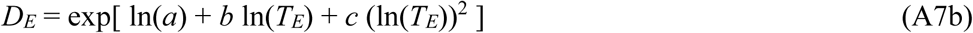

where *T*_*M*_ and *T*_*E*_ are mean temperatures in the metalimnion and epilimnion, respectively, and coefficients ln(*a*) = 3.4, *b* = 0.22, and *c* = −0.3 come from (Bottrell et al. 1976). Mean development time at each lake-date, *D*_*ave*_, is then just the weighted average of *D*_*E*_ and *D*_*M*_:

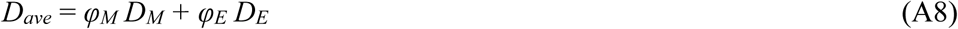

where *φ*_*M*_ and *φ*_*E*_ are the proportion of time per day spent in the metalimnion and epilimnion, respectively (taking into account waning of daylight as autumn progresses).

Then, we calculated birth rate using the egg ratio methods. We calculated the average weighted egg ratio, *E*_*ave*_, using data on infected and uninfected adult host classes. Next we calculated the population-level egg ratio, *E*_*p*_, by multiplying *E*_*ave*_ times the percentage of asexual females in the population. We calculated the per capita birth rate, *b*:

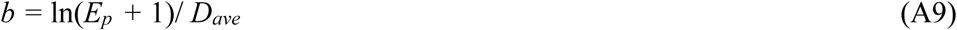

where *D*_*ave*_ follows equ. A8. We calculated the instantaneous rate of increase, *r* = ln (*N*_*t+τ*_ / *N*_*t*_) / *τ*, where *N*_*t+τ*_ / *N*_*t*_ is the ratio of host density between sampling intervals *τ*. Then, death rate during epidemics is *d+pv* = *b* – *r*.

##### Example calculation of index of foraging depression

The ‘index of foraging depression’ becomes more tangible with an illustrative example from one of the 13 studied lakes. As an epidemic unfolded in Goodman Lake, spore yield lagged behind prevalence through time (Fig. A4a), and mean size of uninfected and infected host changed slightly (Fig. A4b). The size-only effect had a modest influence on mean foraging of adult hosts (‘only size’ solid lines in Fig. A4c, as parameterized for the three laboratory-assayed genotypes [Table A1]). However, foraging dropped considerably once infection was modeled (‘spore-depressed’ dashed lines in Fig. A4c). The index of foraging depression comes from the difference between lines ‘only size’ and ‘spore-depressed’ lines (Fig. A4d). The index of foraging depression, *FD*, for a given lake-date-genotype combination is:

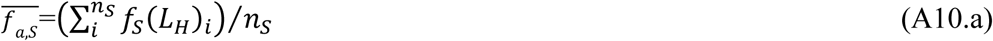

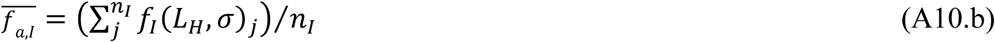

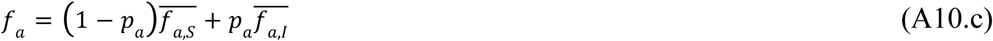

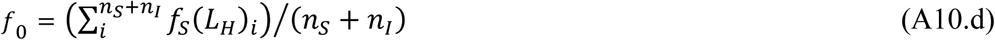

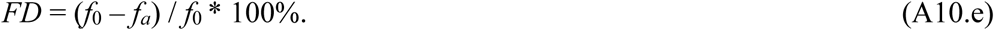

Here, mean feeding rate of the sample of susceptible (uninfected) adults, 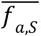, is calculated for *n*_*S*_ individuals using foraging function *f*_*S*_(*L*_*H*_) (equ. 2.a) for individual *i* given its body length *L*_*H*_ (equ. A10.a). Similarly, mean feeding rate of infected adults, 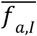, calculated for *n*_*I*_ individuals using foraging function *f*_*I*_(*L*_*H*_,*σ*) (equ. 2.b) for individual *j* given its body length *L*_*H*_ and the mean spore yield per infected host on that sampling date, *σ* (equ. A10.b). The mean foraging rate, *f*_*a*_, is then the average of these mean rates weighted by prevalence of infection in adults, *p*_*a*_ (equ. A10.c). For comparison, we calculated mean adult foraging rate assuming only length (*L*_*H*_) influenced it, *f*0 (equ. A10.d). Specifically, to calculate *f*0 we used the foraging function for susceptibles, *f*_*S*_(*L*_*H*_) (equ. 2a) for each susceptible and infected individual *i* (summed over the total sample, *n*_*S*_ + *n*_*I*_). (This calculation therefore assumed size-specific foraging rate for infected hosts, 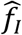, was set to that of susceptible hosts, 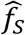, and that spore accrual did not suppress feeding, so *α* = 0; note that this is equivalent to the *‘size only*’ model 2a from Table 1). The index of foraging depression, *FD*, was then the relative depression of foraging due to disease (equ. A10.e). In a given lake, we calculated three separate values of *FD* each sampling date: one for each set of genotype-specific parameter estimates from the foraging rate experiment (Table A1; these produced the three lines in Fig. A4.d). We calculated the temporal mean for each of the three genotype-specific *FD* values, and then averaged across those three temporal means to produce one value of *FD* per lake (plotted in Fig. 5b,c).

**Table A1.** Statistical description of the winning foraging function, ‘5b: *size and spores, linear*’ (from Table 1; also presented as equ. 2). (a) Best-fit parameter estimates (with bootstrapped lower and upper 95% CI) for size-specific foraging rates of uninfected hosts 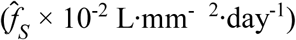 and infected hosts 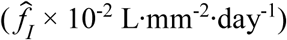, and for the linear sensitivity coefficient (α × 10^−5^ mm^3^·spore^-1^). (b) *P*-values of permutation tests comparing those parameter estimates between host genotypes (9,999 randomizations per contrast; asterisks indicate significant pairwise differences after Holm–Bonferroni correction). These parameters generate the curves shown in the text (Fig. 1) and were also paired with field data in the calculation of the index of foraging depression (Fig. A4; Fig. 5b,c).

| (a) Estimate (95% CI) for each host genotype |  |  |  |
| --- | --- | --- | --- |
| Parameter | A4-4 | BD-30 | STD |
| $\hat{f}_S$ | 1.94 (1.83, 2.05) | 1.78 (1.62, 1.96) | 1.69 (1.53, 1.85) |
| $\hat{f}_I$ | 1.54 (1.41, 1.67) | 1.31 (1.17, 1.46) | 0.90 (0.80, 1.00) |
| $\alpha$ | 2.49 (2.18, 2.85) | 2.86 (2.23, 3.35) | 0.92 (0.22, 1.59) |
| (b) *P*-value for comparison between genotypes |  |  |  |
| Parameter | A4-4 vs. BD-30 | BD-30 vs. STD | STD vs. A4-4 |
| $\hat{f}_S$ | 0.028 | 0.26 | 0.0002 * |
| $\hat{f}_I$ | 0.0017 * | < 0.0001 * | < 0.0001 * |
| $\alpha$ | 0.095 | < 0.0001 * | < 0.0001 * |

### APPENDIX FIGURE LEGENDS

**Figure A1.**
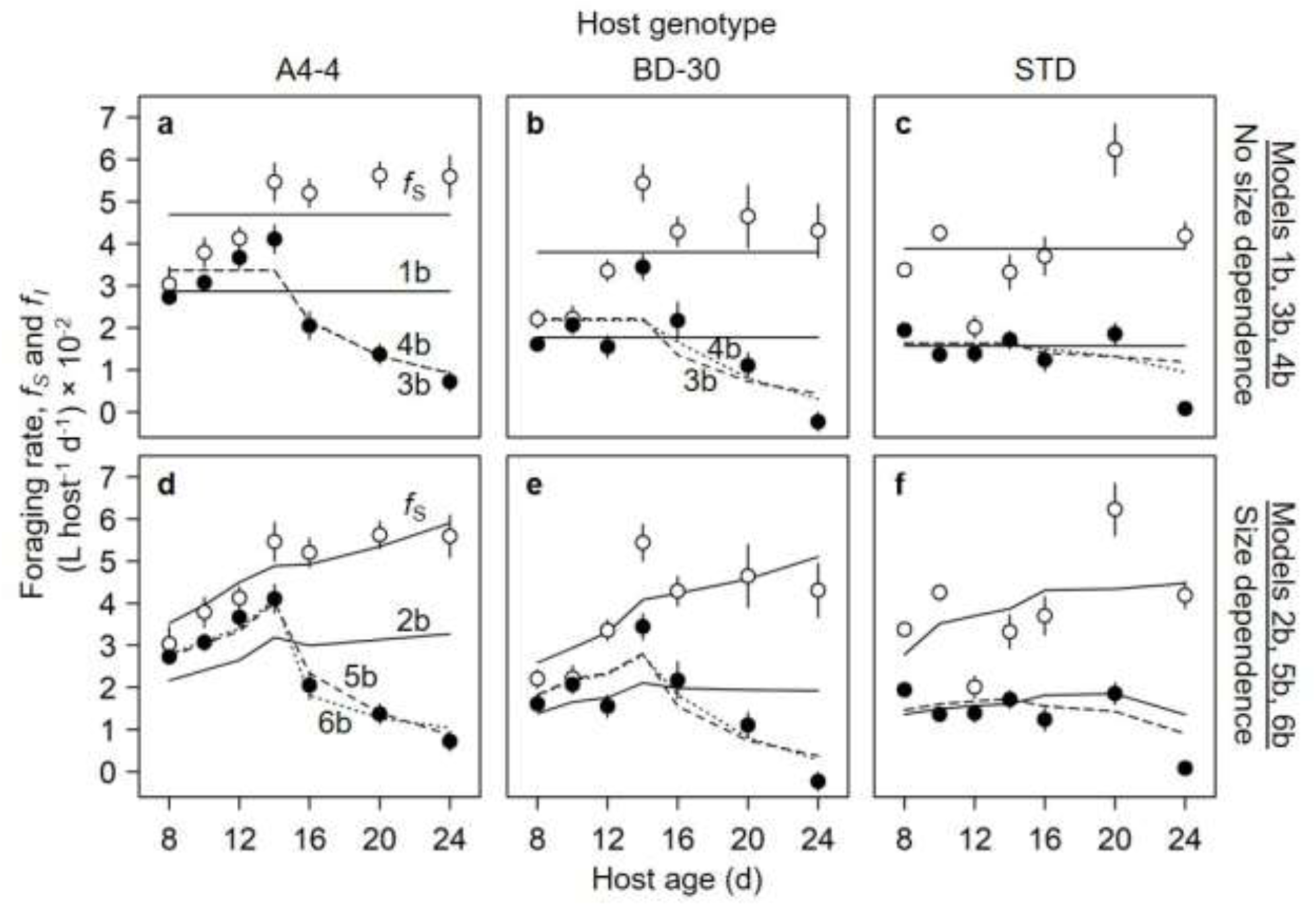
Functions of spore-dependent foraging rate, *f*(***L***_***H***_,*σ*), in models 1b-6b (see Table 1). These functions were fit to observed algal data for uninfected (***f***_***S***_, white) and infected (***f***_***I***_, black) hosts from three genotypes but visualized with calculated foraging rate (mean ± 1 SE). (a–c) *No size dependence*: Foraging functions 1b (solid line), 3b (dashed), and model 4b (dotted), respectively, do not scale with host surface area (***L***_***H***_); in panel a, 3b and 4b overlap. (d-f) *Size dependence*: Function 2b (solid), 5b (dashed; this function is equ. 2 and also plotted in Fig. 1a-c), and 6b (dotted) depend on surface area; in panel f, 5b and 6b overlap. Note that for uninfected hosts with *σ*=0, models 1b, 3b, and 4b are equivalent and models 2b, 5b, and 6b are equivalent (***f***_***S***_, solid). Genotypes: A4-4 (panels a,d), Beaver Dam-30 (‘BD-30’; b,e), and standard (‘STD’; c,f).

**Figure A2.**
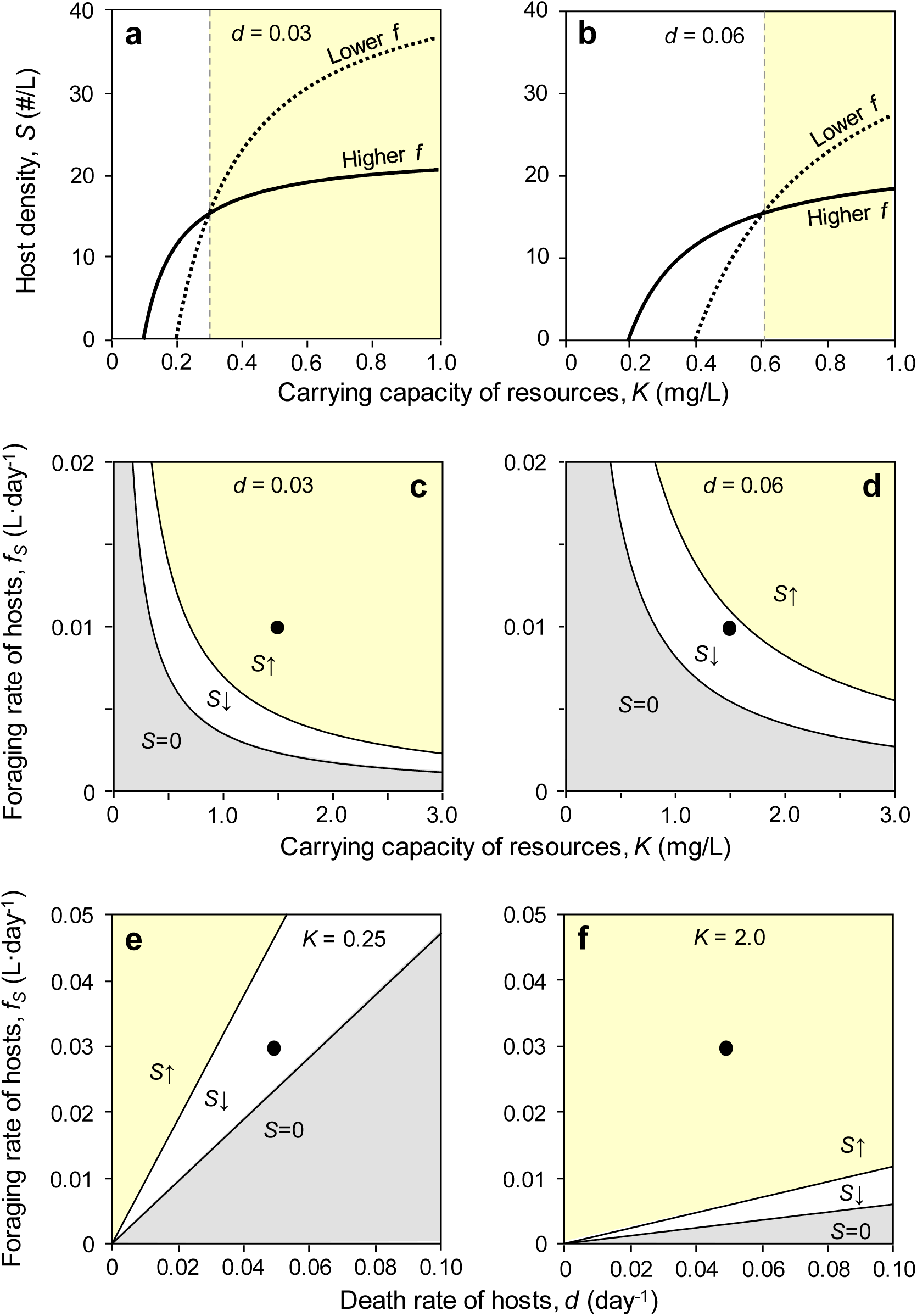
Graphical response of hosts (*S*^*^) to depressed foraging rate (*f*) without disease. (a,b) Host density at lower *f* (dashed; 0.0175 L·day^-1^) and higher *f* (solid; 0.0350 L·day^-1^). In the yellow region, hosts with lower *f* are more abundant, illustrated for (a) lower mortality (*d*=0.03 day^-1^) and (b) higher mortality (*d*=0.06 day^-1^). (c)-(d) Regions of carrying capacity (*K*) and feeding rate (*f*) in which hosts increase with lower feeding rate (yellow; ‘S↑’; -∂S*/∂f > 0; equ. A4.a), hosts decrease with lower feeding rate (white; ‘S↓’; -∂S*/∂f < 0), or hosts cannot persist (‘*S*=0’; *K* < *A*^*^), for (c) lower mortality (*d*=0.03 day^-1^) and (d) higher mortality (*d*=0.06 day^-1^). (e)-(f) Regions of death rate (d)-feeing rate (f) parameter space supporting those same three states (S↑, S↓, *S*=0). Other parameters follow Table 2. (Dots in panels c-f are referred to in text).

**Figure A3.**
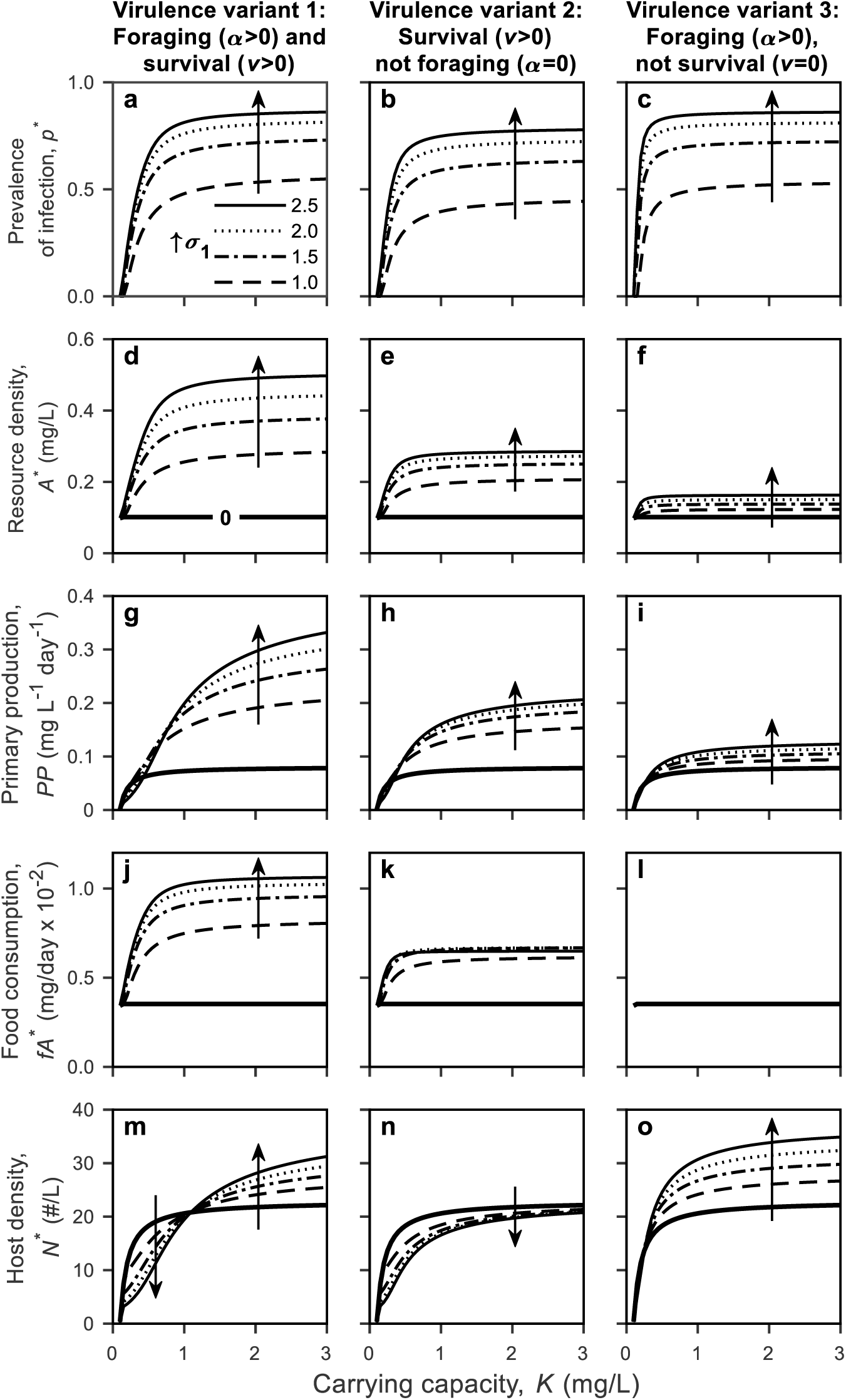
More results from the dynamical model, simulated under ‘virulence variants’. Left column: both foraging depression and virulence on survival (*α* > 0, *v* > 0; shown in Fig. 2). Middle column: only virulence on survival (*α* = 0, *v* > 0). Right column: only foraging depression (*α* > 0, *v* = 0). (a-c) Equilibrial prevalence of infection, *p*^*^. (d–f) Resource density, *A*^*^. (g–i) Primary production, *PP*= *r*_*m*_ *A*^*^(1 - *A*^*^/*K*). (j-l) Food consumption per host, *fA*^*^. (m-o) Total host density, *N*^*^ = *S*^*^ + *I*^*^. Arrows point along contours of increasing maximum spore yields, *σ*_1_. Disease-free conditions (‘0’) noted with thick solid contours. Parameters follow Table 2.

**Figure A4.**
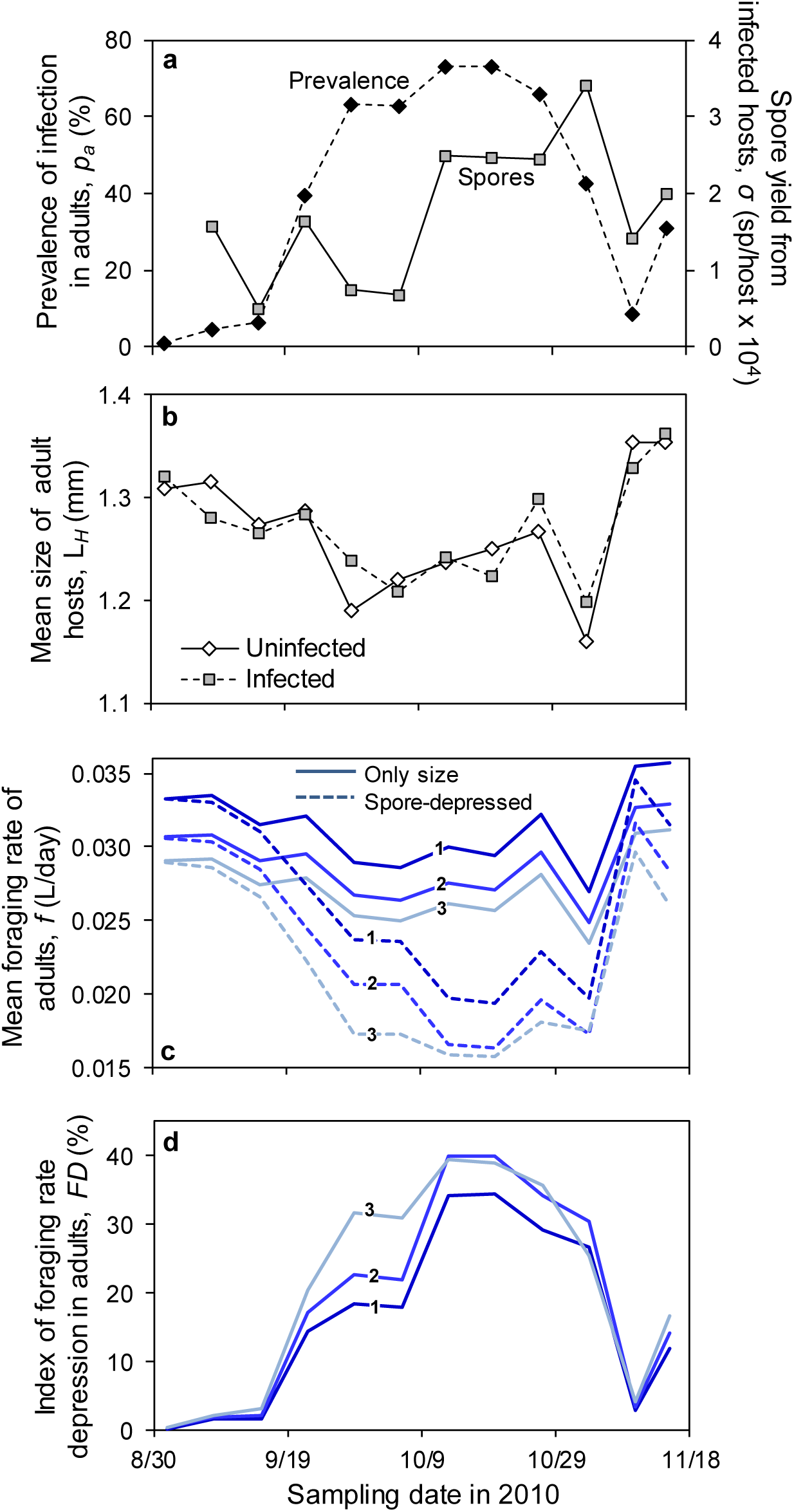
Illustration of the ‘foraging depression’ index, calculated with adult hosts (shown in Fig. 5). (a) An example of a large fungal epidemic in Goodman Lake (Fig. 4a,c): prevalence of infection of adults (percentage infected; black diamonds) and spore yield per infected host (*σ*, grey squares). (b) Mean size of infected and uninfected adults (*L*_*H*_). (c) Components of the foraging rate (*f*) depression index (equ. A10), calculated for clonal genotypes 1 (A4-4), 2 (BD-30), and 3 (STD; parameters in Table A1). The ‘only size’ lines (solid) calculate foraging rate based on host size alone. ‘Spore-depressed’ lines (dashed) assume different size-specific foraging rates for infected 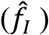 and uninfected 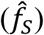 hosts, and spore-mediated foraging depression (proportional to α). (d) Percentage decrease from the ‘only size’ to ‘spore-depressed’ estimates. This calculation shows that spore accumulation within hosts strongly depresses mean foraging rate of adults in this population of *Daphnia*.

## References

Abrams, P. A. 2009. When does greater mortality increase population size? The long history and diverse mechanisms underlying the hydra effect. Ecology Letters 12:462–474.

Abrams, P. A., and H. Matsuda. 2005. The effect of adaptive change in the prey on the dynamics of an exploited predator population. Canadian Journal of Fisheries and Aquatic Sciences 62:758–766.

Altizer, S., R. S. Ostfeld, P. T. J. Johnson, S. Kutz, and C. D. Harvell. 2013. Climate change and infectious diseases: from evidence to a predictive framework. Science 341:514–519.

Anagnostakis, S. L. 1982. Biological control of chestnut blight. Science 215:466–471.

Anderson, R. M., and R. M. May. 1979. Population biology of infectious diseases: Part I. Nature 280:361–367.

Anderson, R. M., and R. M. May. 1981. The population dynamics of microparasites and their invertebrate hosts. Philosophical Transactions of the Royal Society of London. B, Biological Sciences 291:451–524.

Auld, S. K. J. R., S. R. Hall, and M. A. Duffy. 2012. Epidemiology of a *Daphnia*-multiparasite system and its implications for the Red Queen. Plos One 7:6.

Auld, S. K. J. R., S. R. Hall, J. H. Ochs, M. Sebastian, and M. A. Duffy. 2014. Predators and patterns of within-host growth can mediate both among-host competition and evolution of transmission potential of parasites. American Naturalist 184:S77–S90.

Auld, S. K. J. R., R. M. Penczykowski, J. H. Ochs, D. C. Grippi, S. R. Hall, and M. A. Duffy. 2013. Variation in costs of parasite resistance among natural host populations. Journal of Evolutionary Biology 26:2479–2486.

Boots, M., A. Best, M. R. Miller, and A. White. 2009. The role of ecological feedbacks in the evolution of host defence: what does theory tell us? Philosophical Transactions of the Royal Society B-Biological Sciences 364:27–36.

Borer, E. T., E. W. Seabloom, J. B. Shurin, K. E. Anderson, C. A. Blanchette, B. Broitman, S. D. Cooper et al. 2005. What determines the strength of a trophic cascade? Ecology 86:528–537.

Bottrell, H. H., A. Duncan, Z. M. Gliwicz, E. Grygierek, A. Herzig, A. Hillbricht Ilkowska, H. Kurasawa et al. 1976. A review of some problems in zooplankton production studies.

Buck, J. C., and W. J. Ripple. 2017. Infectious agents trigger trophic cascades. Trends in Ecology & Evolution 32:681–694.

Burnham, K. P., and D. R. Anderson. 2002, Model selection and multimodel inference: a practical information-theoretic approach, 2nd ed. New York, Springer-Verlag.

Case, T. J. 2000, An Illustrated Guide to Theoretical Ecology. New York, Oxford University Press.

Chen, C. T., and F. J. Millero. 1977. Use and misuse of pure water PVT properties for lake waters. Nature 266:707–708.

Civitello, D. J., R. M. Penczykowski, A. N. Smith, M. S. Shocket, M. A. Duffy, and S. R. Hall. 2015. Resources, key traits and the size of fungal epidemics in *Daphnia* populations. Journal of Animal Ecology 84:1010–1017.

Cleaveland, S., M. K. Laurenson, and L. H. Taylor. 2001. Diseases of humans and their domestic mammals: pathogen characteristics, host range and the risk of emergence. Philosophical Transactions of the Royal Society B-Biological Sciences 356:991–999.

Cooper, J., R. J. M. Crawford, M. S. De Villiers, B. M. Dyer, G. J. G. Hofmeyr, and A. Jonker. 2009. Disease outbreaks among penguins at sub-Antarctic Marion Island: a conservation concern Marine Ornithology 37:193–196

Cortez, M. H., and P. A. Abrams. 2016. Hydra effects in stable communities and their implications for system dynamics. Ecology 97:1135–1145.

Daszak, P., L. Berger, A. A. Cunningham, A. D. Hyatt, D. E. Green, and R. Speare. 1999. Emerging infectious diseases and amphibian population declines. Emerging Infectious Diseases 5:735–748.

de Roos, A. M., and L. Persson. 2013, Population and Community Ecology of Ontogenetic Development. Princeton, New Jersey, USA, Princeton University Press.

Duffy, M. A. 2007. Selective predation, parasitism, and trophic cascades in a bluegill-*Daphnia*-parasite system. Oecologia 153:453–460.

Duffy, M. A., and S. E. Forde. 2009. Ecological feedbacks and the evolution of resistance. Journal of Animal Ecology 78:1106–1112.

Duffy, M. A., and S. R. Hall. 2008. Selective predation and rapid evolution can jointly dampen effects of virulent parasites on *Daphnia* populations. American Naturalist 171:499–510.

Duffy, M. A., S. R. Hall, A. J. Tessier, and M. Huebner. 2005. Selective predators and their parasitized prey: Are epidemics in zooplankton under top-down control? Limnology and Oceanography 50:412–420.

Dwyer, G., and J. S. Elkinton. 1993. Using simple models to predict virus epizootics in gypsy moth populations. Journal of Animal Ecology 62:1–11.

Ebert, D. 2005, Ecology, Epidemiology and Evolution of Parasitism in Daphnia. Bethesda, MD, National Library of Medicine (US), National Center for Biotechnology Information.

Frick, W. F., J. F. Pollock, A. C. Hicks, K. E. Langwig, D. S. Reynolds, G. G. Turner, C. M. Butchkoski et al. 2010. An emerging disease causes regional population collapse of a common North American bat species. Science 329:679–682.

Fry, W. E., and S. B. Goodwin. 1997. Re-emergence of potato and tomato late blight in the United States. Plant Disease 81:1349–1357.

Gotelli, N. J., and A. M. Ellison. 2004, A Primer of Ecological Statistics. Sunderland, MA, Sinauer Associates, Inc.

Green, J. 1974. Parasites and epibionts of *Cladocera*. Transactions of the Zoological Society of London 32:417–515.

Grover, J. P. 1995. Competition, herbivory, and enrichment: nutrient-based models for edible and inedible plants. The American Naturalist 145:746–774.

Grover, J. P. 1997, Resource Competition: Population and Community Biology Series. Boston, MA, Springer.

Hall, S. R., C. Becker, and C. E. Cáceres. 2007a. Parasitic castration: a perspective from a model of dynamic energy budgets. Integrative and Comparative Biology 47:295–309.

Hall, S. R., C. R. Becker, M. A. Duffy, and C. E. Cáceres. 2010. Variation in resource acquisition and use among host clones creates key epidemiological trade-offs. American Naturalist 176:557–565.

Hall, S. R. 2011. Epidemic size determines population-level effects of fungal parasites on *Daphnia* hosts. Oecologia 166:833–842.

Hall, S. R., C. R. Becker, J. L. Simonis, M. A. Duffy, A. J. Tessier, and C. E. Cáceres. 2009a. Friendly competition: evidence for a dilution effect among competitors in a planktonic host-parasite system. Ecology 90:791–801.

Hall, S. R., M. A. Duffy, A. J. Tessier, and C. E. Cáceres. 2005. Spatial heterogeneity of daphniid parasitism within lakes. Oecologia 143:635–644.

Hall, S. R., J. L. Simonis, R. M. Nisbet, A. J. Tessier, and C. E. Cáceres. 2009b. Resource ecology of virulence in a planktonic host-parasite system: an explanation using dynamic energy budgets. American Naturalist 174:149–162.

Hall, S. R., L. Sivars-Becker, C. Becker, M. A. Duffy, A. J. Tessier, and C. E. Cáceres. 2007b. Eating yourself sick: transmission of disease as a function of foraging ecology. Ecology Letters 10:207–218.

Hilker, F. M., M. Langlais, and H. Malchow. 2009. The Allee effect and infectious diseases: extinction, multistability, and the (dis-)appearance of oscillations. American Naturalist 173:72–88.

Hite, J. L., and C. E. Cressler. 2019. Parasite-mediated anorexia and nutrition modulate virulence evolution. Integrative and Comparative Biology 59:1264–1274.

Hite, J. L., R. M. Penczykowski, M. S. Shocket, K. A. Griebel, A. T. Strauss, M. A. Duffy, C. E. Cáceres et al. 2017. Allocation, not male resistance, increases male frequency during epidemics: a case study in facultatively sexual hosts. Ecology 98:2773–2783.

Hite, J. L., A. C. Pfenning, and C. E. Cressler. 2020. Starving the enemy? Feeding behavior shapes host-parasite interactions. Trends in Ecology & Evolution 35:68–80.

Hochachka, W. M., and A. A. Dhondt. 2000. Density-dependent decline of host abundance resulting from a new infectious disease. Proceedings of the National Academy of Sciences of the United States of America 97:5303–5306.

Hurtado, P. J., S. R. Hall, and S. P. Ellner. 2014. Infectious disease in consumer populations: dynamic consequences of resource-mediated transmission and infectiousness. Theoretical Ecology 7:163–179.

Johnson, P. T. J., A. R. Townsend, C. C. Cleveland, P. M. Glibert, R. W. Howarth, V. J. McKenzie, E. Rejmankova et al. 2010. Linking environmental nutrient enrichment and disease emergence in humans and wildlife. Ecological Applications 20:16–29.

Klüttgen, B., U. Dulmer, M. Engels, and H. T. Ratte. 1994. ADaM, an artificial freshwater for the culture of zooplankton. Water Research 28:743–746.

Kooijman, S. A. L. M. 2010, Dynamic Energy Budget Theory for Metabolic Organisation. Great Britain, Cambridge University Press.

Lafferty, K. D., and A. M. Kuris. 2009. Parasitic castration: the evolution and ecology of body snatchers. Trends in Parasitology 25:564–572.

Lessios, H. A., D. R. Robertson, and J. D. Cubit. 1984. Spread of *Diadema* mass mortality through the Caribbean. Science 226:335–337.

McIntire, K. M., and S. A. Juliano. 2018. How can mortality increase population size? A test of two mechanistic hypotheses. Ecology 99:1660–1670.

Overholt, E. P., S. R. Hall, C. E. Williamson, C. K. Meikle, M. A. Duffy, and C. E. Cáceres. 2012. Solar radiation decreases parasitism in *Daphnia*. Ecology Letters 15:47–54.

Peacor, S. D., and E. E. Werner. 2001. The contribution of trait-mediated indirect effects to the net effects of a predator. Proceedings of the National Academy of Sciences of the United States of America 98:3904–3908.

Piñeiro, G., S. Perelman, J. P. Guerschman, and J. M. Paruelo. 2008. How to evaluate models: Observed vs. predicted or predicted vs. observed? Ecological Modelling 216:316–322.

Polis, G. A., W. B. Anderson, and R. D. Holt. 1997. Toward an integration of landscape and food web ecology: The dynamics of spatially subsidized food webs. Annual Review of Ecology and Systematics 28:289–316.

Preston, D. L., and E. L. Sauer. 2020. Infection pathology and competition mediate host biomass overcompensation from disease. Ecology 101.

Roelke-Parker, M. E., L. Munson, C. Packer, R. Kock, S. Cleaveland, M. Carpenter, S. J. Obrien et al. 1996. A canine distemper virus epidemic in Serengeti lions (*Panthera leo*). Nature 379:441–445.

Sanderson, C. E., and K. A. Alexander. 2020. Unchartered waters: Climate change likely to intensify infectious disease outbreaks causing mass mortality events in marine mammals. Global Change Biology. In press.

Sarnelle, O., and A. E. Wilson. 2008. Type III functional response in *Daphnia*. Ecology 89:1723–1732.

Schröder, A., L. Persson, and A. M. de Roos. 2009. Culling experiments demonstrate size-class specific biomass increases with mortality. Proceedings of the National Academy of Sciences of the United States of America 106:2671–2676.

Shocket, M. S., A. T. Strauss, J. L. Hite, M. Sljivar, D. J. Civitello, M. A. Duffy, C. E. Cáceres et al. 2018. Temperature drives epidemics in a zooplankton-fungus disease system: a trait-driven approach points to transmission via host foraging. American Naturalist 191:435–451.

Shurin, J. B., and E. W. Seabloom. 2005. The strength of trophic cascades across ecosystems: predictions from allometry and energetics. Journal of Animal Ecology 74:1029–1038.

Stewart Merrill, T. E., and C. E. Cáceres. 2018. Within-host complexity of a plankton-parasite interaction. Ecology 99:2864–2867.

Stewart Merrill, T. E., S. R. Hall, L. Merrill, and C. E. Cáceres. 2019. Variation in immune defense shapes disease outcomes in laboratory and wild *Daphnia*. Integrative and Comparative Biology 59:1203–1219.

Strauss, A. T., A. M. Bowling, M. A. Duffy, C. E. Cáceres, and S. R. Hall. 2018. Linking host traits, interactions with competitors and disease: Mechanistic foundations for disease dilution. Functional Ecology 32:1271–1279.

Strauss, A. T., D. J. Civitello, C. E. Cáceres, and S. R. Hall. 2015. Success, failure and ambiguity of the dilution effect among competitors. Ecology Letters 18:916–926.

Strauss, A. T., J. L. Hite, D. J. Civitello, M. S. Shocket, C. E. Cáceres, and S. R. Hall. 2019. Genotypic variation in parasite avoidance behaviour and other mechanistic, nonlinear components of transmission. Proceedings of the Royal Society B-Biological Sciences 286.

Tessier, A. J., and P. Woodruff. 2002. Cryptic trophic cascade along a gradient of lake size. Ecology 83:1263–1270.

Vredenburg, V. T., R. A. Knapp, T. S. Tunstall, and C. J. Briggs. 2010. Dynamics of an emerging disease drive large-scale amphibian population extinctions. Proceedings of the National Academy of Sciences of the United States of America 107:9689–9694.

Webb, D. J., B. K. Burnison, A. M. Trimbee, and E. E. Prepas. 1992. Comparison of chlorophyll *a* extractions with ethanol and dimethyl sulfoxide/acetone, and a concern about spectrophotometric phaeopigment correction. Canadian Journal of Fisheries and Aquatic Sciences 49:2331–2336.

Welschmeyer, N. A. 1994. Fluorometric analysis of chlorophyll *a* in the presence of chlorophyll *b* and pheopigments. Limnology and Oceanography 39:1985–1992.

